# Endosomal Membrane Tension Regulates ESCRT-III-Dependent Intra-Lumenal Vesicle Formation

**DOI:** 10.1101/550483

**Authors:** Vincent Mercier, Jorge Larios, Guillaume Molinard, Antoine Goujon, Stefan Matile, Jean Gruenberg, Aurélien Roux

**Affiliations:** Department of Biochemistry, University of Geneva, CH-1211 Geneva, Switzerland; National Center of Competence in Research Chemical Biology, University of Geneva, CH-1211 Geneva, Switzerland; Department of Organic Chemistry, University of Geneva, CH-1211 Geneva, Switzerland

## Abstract

Plasma membrane tension strongly affects cell surface processes, such as migration, endocytosis and signalling. However, it is not known whether membrane tension of organelles regulates their functions, notably intracellular traffic. The ESCRT-III complex is the major membrane remodelling complex that drives Intra-Lumenal Vesicle (ILV) formation on endosomal membranes. Here, we made use of a new fluorescent membrane tension probe to show that ESCRT-III subunits are recruited onto endosomal membranes when membrane tension is reduced. We find that tension-dependent recruitment is associated with ESCRT-III polymerization and membrane deformation in vitro, and correlates with increased ILVs formation in ESCRT-III decorated endosomes in vivo. Finally, we find that endosomal membrane tension decreases when ILV formation is triggered by EGF under physiological conditions. These results indicate that membrane tension is a major regulator of ILV formation and of endosome trafficking, leading us to conclude that membrane tension can control organelle functions.

**One Sentence Summary:** Membrane tension decrease facilitates membrane remodeling by ESCRT-III polymerization during intra-lumenal vesicle formation.

---

Basic cellular functions are controlled by plasma membrane tension ^1^. We wondered whether membrane tension could also regulate the remodelling of endosomal membranes. The ESCRT-III complex functions as a general membrane deformation and fission machinery in an orientation opposite to endocytosis, away from the cytoplasm. It plays essential roles in cytokinetic abscission, viral budding, nuclear envelope reformation ^2–5^, as well as plasma membrane ^6^ and lysosome membrane repair ^7^. As it drives ILV formation in endosomes ^7^, we wondered whether ESCRT-III function could be controlled by endosomal membrane tension.

When cells are bathed in a hypertonic solution, the cytoplasmic volume is reduced through water expulsion, which results in a decrease of plasma membrane tension ^8^. We reasoned that, after hypertonic shock, the endosome volume and in turn membrane tension should also be reduced (see outline Fig S1a). We thus decided to determine whether hypertonic conditions affected ESCRT-III membrane association, and, if so, whether the process depended on membrane tension. To test this, we used HeLa-Kyoto cells stably expressing the ESCRT-III subunit CHMP4B-GFP at low, near-endogenous levels. In these cells, CHMP4B-GFP showed mostly a diffuse, nuclear and cytosolic pattern, with few dots presumably corresponding to endosomes (Fig 1a). This distribution changed dramatically after hyperosmotic shock (+0.5M sucrose, final osmolarity ∼830mOsm): CHMP4B-GFP relocalized to cytoplasmic punctae (Fig 1a, Movie S1), as did endogenous CHMP4B in HeLa MZ cells (Fig S1f). The redistribution was both rapid (Fig 1b-c) and transient (Fig 1d): CHMP4B punctae appeared within minutes after hypertonic shock (Fig 1a-b,d,f), with a maximum at ≈20 min (Fig 1d), and then disappeared with slower kinetics (Fig 1d-e, Movie S2). The number of CHMP4B punctae increased with increasing osmolarity above the physiological level (Fig S1g). By contrast, neither hypotonic nor isotonic medium addition triggered CHMP4B relocalization (Fig 1b and Fig S1d). Relocalization was not dependent on the chemical nature of the osmolyte, as it could be recapitulated with the addition of 250mM NaCl to a final osmolarity ∼830mOsm (Fig S1c). Furthermore, the disappearance of CHMP4B punctae could be accelerated by replacing the hypertonic medium with isotonic medium (Fig S1k) or even more so with hypotonic medium (Fig 1f-g).

**Figure 1:**
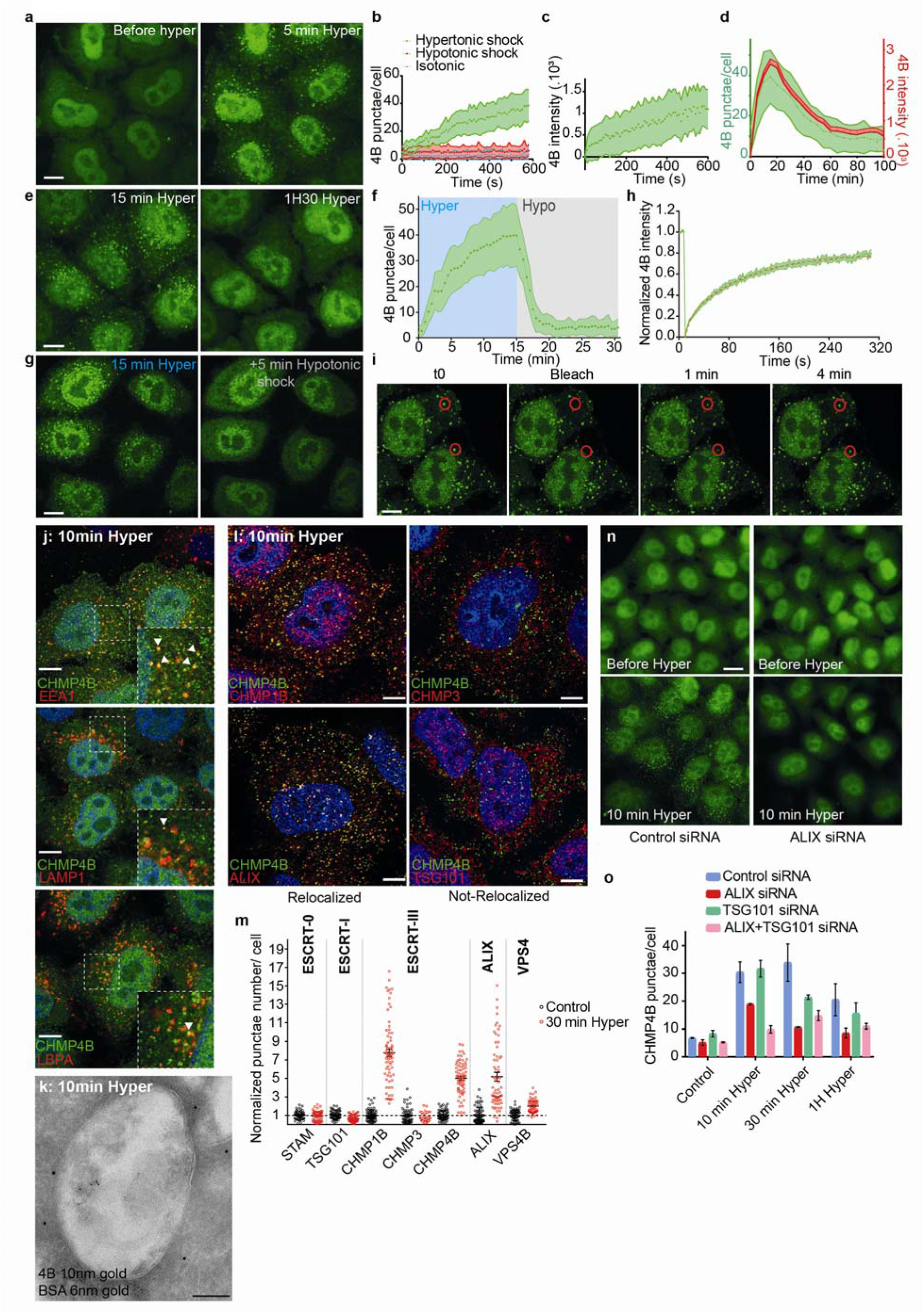
Hypertonic shock triggers fast and transient CHMP4B recruitment on endosomes. a) Confocal projections of HeLa cells stably expressing CHMP4B-GFP before and after a 5min incubation with a hypertonic solution (800 mOsm). Bar: 10 µm. b) Average number of CHMP4B punctae per cell with time after hypertonic (∼800 mOsm), isotonic (∼330 mOsm) or hypotonic (∼250 mOsm) shocks. Shaded areas: mean ± SEM (N=27-31 cells from 3 independent replicates). c) Mean intensity of CHMP4B-GFP punctae over time. Shaded areas: mean ± SD (N=821 punctae from 19 cells from 3 independent replicates). d) Mean intensity of CHMP4B punctae (Red curve, red shaded area: SEM) and average number of CHMP4B punctae per cell (green curve, green shaded area: SD) over 100 min. (N=27 cells, 3 independents replicates). e) Confocal projections of CHMP4B-GFP HeLa cells for later time points (15 and 90 min) after hypertonic shock. Bar: 10 µm. f) Average number of CHMP4B-GFP punctae per cell during a 15min hypertonic shock (∼800 mOsm, Blue) followed by 15min of hypotonic shock (∼250 mOsm, grey). Shaded area: SD (N=47 cells from 3 independent replicates). g) Representative confocal projections of one experiment quantified in (f). Bar: 10 µm. h) Mean fluorescence recovery curve after photobleaching of CHMP4B-GFP punctae, fitted with a double exponential (dotted line). Shaded area: SEM (N=33 punctae from 3 independent replicates). i) Representative confocal images of one experiment quantified in (h). Bar: 5 µm. j) Confocal images of immunofluorescences showing colocalization (arrows) of several endosomal markers with CHMP4B-GFP after 10min hypertonic shock. Bar: 5 µm. k) BSA-gold (6nm) was endocytosed for 15min followed by a 10min hypertonic shock. Cells were processed for immuno-electron microscopy using anti CHMP4B antibody followed by 10nm ProteinA-gold. Bar: 100 nm. l) Confocal images of immunofluorescences showing colocalization between CHMP4B-GFP and indicated ESCRT subunits after 10min hypertonic shock. Bar: 4 µm. m) Quantification of punctae number per cell from automated confocal images of immunofluorescence before (black) and after 30min hypertonic shock (red) for various markers. Bars are mean±SEM. One dot corresponds to the average number of punctae per cell in one field of view (70 fields in 3 independent experiments, few tens of cells per field) normalized to the average number before shock (black). n) Confocal images of CHMP4B-GFP HeLa cells before and after 10min hypertonic shock, pre-treated with anti-ALIX siRNAs or control siRNAs against VSV-G. Bar: 15 µm. o) Mean number of CHMP4B punctae per cell, for cells treated with control siRNAs or siRNAs against ALIX, TSG101 or both, before and after hypertonic shocks of various duration. (N>2000 cells, from 3 independent replicates).

These punctae were not CHMP4B aggregates, as after photobleaching CHMP4B fluorescence recovered to ≈ 80% of the initial value with a t1/2 ≈ 1 min (Fig 1h-i), showing that subunits were readily exchanged with cytosolic CHMP4B. Moreover, this fast turnover was inhibited by overexpression of the dominant negative mutant K173Q of the triple A ATPase VPS4 (Figure S1e). Because Vps4-dependent high-turnover of ESCRT-III subunits is associated with functionality of the ESCRT-III assemblies ^9^, our results support the view that CHMP4B punctae observed after hypertonic shock are functional assemblies of ESCRT-III. We conclude that hypertonic conditions stimulated the rapid and transient recruitment of ESCRT-III onto cytoplasmic structures.

We next wondered whether these structures were endosomes. Indeed, CHMP4B after hypertonic shock colocalized with the early endosomal marker EEA1 (Figure 1j), and to a lesser extent with the late endosomal markers LAMP1 and lysobisphosphatidic acid (LBPA) (Fig 1j). Similarly, CHMP4B colocalized with internalized transferrin (Fig S1i-j) or EGF (Fig S1h) labelling early endosomes — but to a lesser extent with chased EGF labelling late endosomes (Fig S1h). Consistently, subcellular fractionation showed that CHMP4B was increased in endosomal membrane fractions after hypertonic shock (Fig S1b). Finally, immunogold-labelling of cryo-sections showed that, after hypertonic shock, the limiting membrane of endosomes containing BSA-gold endocytosed for 15min was decorated with CHMP4B antibody (Fig 2k, FigS2a). Among other ESCRT-III subunits, CHMP1B and VPS4 showed an enhanced punctate distribution, but not CHMP3 (Fig 1l-m; Fig S2b, Fig S3a). Altogether, these observations show that most ESCRT-III subunits were recruited onto endosomes after hypertonic shock.

**Figure 2:**
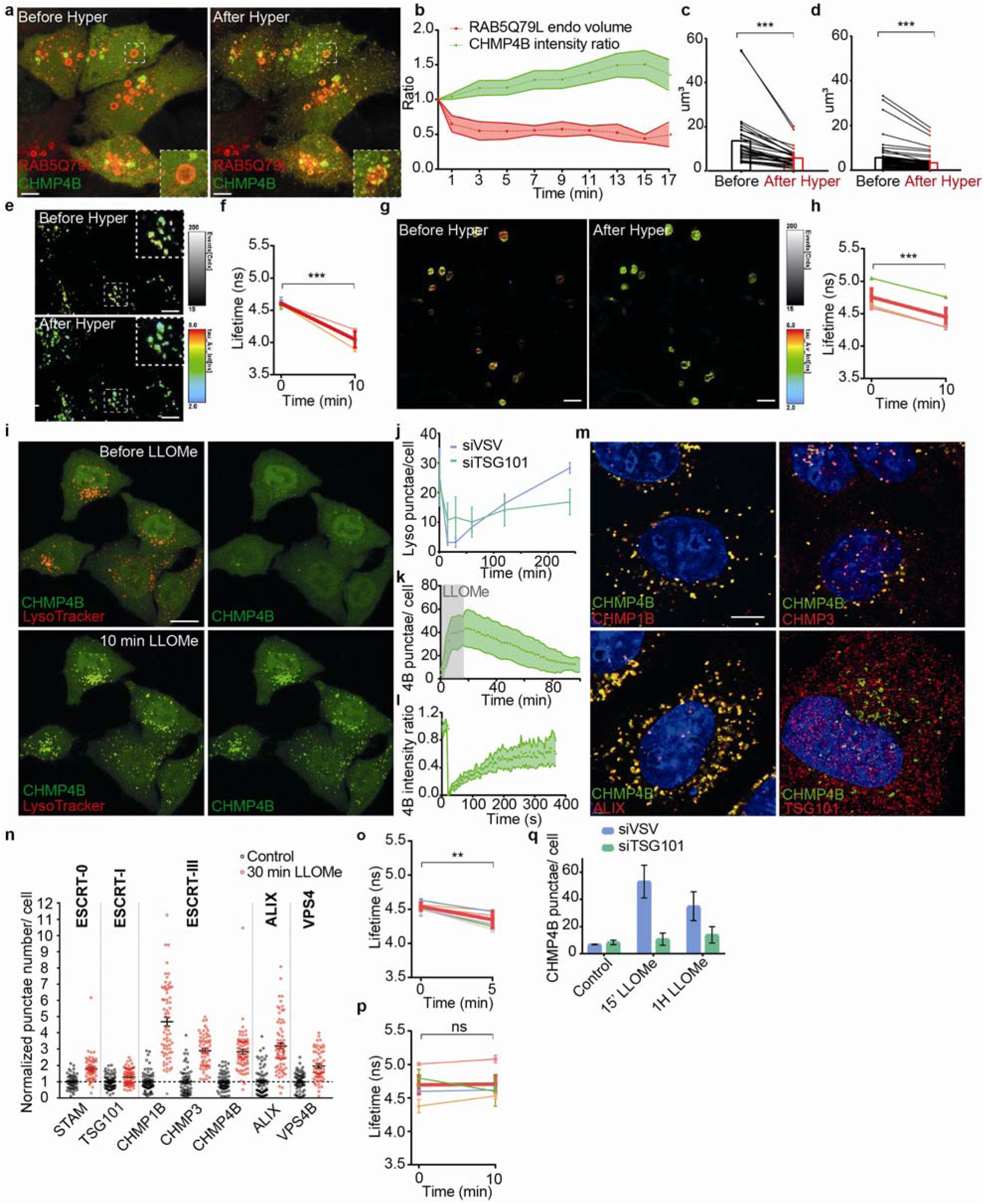
Hypertonic shock and LLOMe decrease membrane tension of endosomes. a) Confocal images of HeLa-CHMP4B-GFP cells expressing mCherry-Rab5Q79L before and after a 20min hypertonic shock. Bar: 5 µm. b) Average volume of Rab5Q79L endosomes (red curve, shaded area is SD) and average intensity of CHMP4B-GFP (green curve, shaded area is SD) after hypertonic shock (time 0), and normalized to initial value. (N=12 RAB5Q79L endosomes). c) Volumes of single RAB5Q79L endosomes before (black) and after a 10min hypertonic shock (red) (N=30 endosomes from 3 independent replicates, two-tailed Wilcoxon test: P=0.0000000019). d) Volumes of single MDA-MB-231 endosomes labeled with FM4-64 before (black) and after (red) a 10min hypertonic shock. (N=52 endosomes from 3 independent replicates, two-tailed Wilcoxon test: P<10^-15^). e) FLIM images of HeLa endosomes labelled with Lyso Flipper before and after 10min hypertonic shock. Bar: 10 µm. f) Lyso Flipper lifetime measurements from (e) before and after hypertonic shock. Thin coloured lines: 5 independents experiments; thick red line: mean with SEM (two-tailed paired t-test: P=0.00000884003). g) FLIM images of FliptR-labelled mCherry-RAB5Q79L endosomes in live HeLa MZ cells before and after a 10min hypertonic shock. Bar: 5 µm. h) FliptR lifetime measurements from 3 independent experiments as shown in (g). Thin coloured lines: single experiments; thick red line: mean±SEM of the 4 experiments (two-tailed paired t-test: P=0.0003338778). i) Confocal images of live HeLa-CHMP4B-GFP cells labelled with LysoTracker before and after a 10min incubation with 0.5mM LLOMe. Bar: 10 µm.j) Average number of LysoTracker punctae per cell (>1000 cells in 3 independent replicates), and for cells treated with control siRNAs (VSV, blue) or siRNAs against TSG101 (green). Error bars: SEM. k) Average number of CHMP4B-GFP punctae per cell with time, during and after LLOMe treatment. Shaded area: SEM (N=33 cells from 3 independent replicates). l) Recovery after photobleaching of individual CHMP4B-GFP punctae induced by LLOMe treatment. Shaded area: SEM. (N=14 endosomes from 3 independent replicates). m) Confocal images of immuno-fluorescence against several markers after 10min incubation with LLOMe treatment. Bar: 4 µm. n) Number of punctae per cell for different ESCRT subunits before (black) and after LLOMe treatment (red). Each dot represents the mean number of ESCRT punctae/cell in one field of view (70 fields in 3 independent experiments, few tens of cells per field) (mean ± SEM). o) Lyso Flipper lifetime measurements before and 5min after LLOMe addition (quantification as in f). Thin coloured lines: 6 independent experiments; thick red line: mean with SEM (two-tailed paired t-test: P=0.00454651). p) FliptR lifetime measurements of RAB5Q79L endosomes before and 10min after LLOMe treatment (lifetime was measured as in f). Thin coloured lines: independents experiments; thick red line: mean with SEM (two-tailed paired t-test: P=0.89443). q) Average number of CHMP4B-GFP punctae per cell before (control) and after treatment with LLOMe in cells transfected with control siRNAs (siVSV) or siRNAs against TSG101. Error bars are SEM (N>1000 cells, from 3 independent replicates). In f, h, o and p, for each experiment, average fluorescent lifetimes were calculated from >500 endosomes taken from at least 3 different cells.

The major components of ESCRT-0 and ESCRT-I, STAM and TSG101 respectively, were not recruited suggesting that those complexes remained unaffected upon hypertonic shock (Fig 1m, Fig S2b, Fig S3a). As ESCRT-0, -I and -II promote ESCRT-III nucleation, we wondered if the ESCRT-III relocalization depended on its known nucleators, ESCRTs and ALIX. ALIX was efficiently recruited onto endosomes after hypertonic shock (Figure 1lm, Fig S2b, Fig S3a). Remarkably, ALIX depletion with siRNAs partially inhibited CHMP4B membrane recruitment (Fig 1n-o; Fig S4), showing that the process depends on ALIX at least in part. While TSG101 depletion only slightly reduced CHMP4B recruitment, the double ALIX-TSG101 knock-down most efficiently inhibited CHMP4B membrane recruitment (Fig 1o; Fig S4), consistent with the view that ALIX and TSG101 function in parallel pathways ^10^. Altogether, these data indicate that hypertonic conditions cause the selective recruitment of ESCRT-III subunits onto endosomal membranes through its nucleators.

Next, we investigated whether hypertonic conditions would also reduce the endosome volume and in turn membrane tension (Fig S1a). In order to measure the volume of individual endosomes, cells were transfected with the constitutively active RAB5 mutant RAB5Q79L, to generate enlarged early endosomes amenable to software-based segmentation. Upon hypertonic shock, CHMP4B was rapidly recruited onto defined regions of these large endosomes (Fig 2a-b), corresponding to ESCRT domains ^11,12^. Strikingly, hypertonic conditions decreased the volume of RAB5Q79L endosomes by more than 50% (Fig 2b-c). Note that the volume change occurs with faster kinetics than the accumulation of CHMP4B (Fig 2a-b), which reflects both the nucleation and polymerization processes. The endosomal volume also decreased by almost 40% after hypertonic shock in non-transfected MDA-MB-231 cells, which have intrinsically large endosomes (Fig 2d). Changes in endosomal volumes are highly unlikely to result from some alterations in the endocytic membrane flux, since hypertonic conditions are known to stop endocytic membrane transport ^13,14^.

We then investigated whether the decrease in endosomal volume observed after hypertonic shock was correlated with a decrease in membrane tension. To this end, we used a modified version of FliptR (Fluorescent lipid tension Reporter, also called Flipper-TR), a probe that reports changes in membrane tension by changes in fluorescence lifetime ^8^, called Lyso Flipper which selectively targets acidic compartments ^15^. After hypertonic shock, the fluorescence lifetime of Lyso Flipper decreased, showing that hypertonicity reduced membrane tension of endosomes (HeLa MZ cells: Fig 2e-f; MDA-MD-231 cells: Fig S5a). This decrease in membrane tension was also observed in cells expressing RAB5Q79L incubated for 2h at 37°C with FliptR in order to label endosomes (Lyso Flipper did not stain well mildly acidic RAB5Q79L endosomes). Indeed, after hypertonic shock, the lifetime of FliptR decreased (Fig 2g-h), and this decrease correlated well with the observed decrease in endosomal volume (Fig 2a-c). These observations suggest that hypertonicity reduced membrane tension of endosomes by deflating them, which in turn may trigger ESCRT-III recruitment. Alternatively, hypertonic conditions may also trigger ESCRT-III recruitment by increasing the cytosolic concentration of its subunits. We thus investigated conditions in which membrane tension could be decreased without changing cytosolic concentrations.

We reasoned that tension may also decrease upon membrane damage. To this end, we used the small peptide LLOMe (L-Leucyl-L-Leucine methyl ester), which causes transient permeabilization of late endosomes and lysosomes ^16^ as illustrated by the rapid neutralization of the endo-lysosomal pH followed by a slower recovery, almost complete after 2h (Fig 2i-j, Fig S6a,c,f-g, Movie S3). Treatment with LLOMe caused fast recruitment of CHMP4B-GFP (Fig2i, k-l, Fig S6b-d,f, Movie S3, S4) onto late endocytic compartments — much like hypertonic conditions (compare Fig 1f with Fig 2k). The effects of LLOMe were transient and followed by re-acidification (Fig2i and Fig S6c), in good agreement with reports ^7,17^ that ESCRT-III-mediated lysosome repair precedes lysophagy and promotes cell survival ^17^.

As observed under hypertonic conditions, CHMP1B, ALIX and VPS4B are recruited onto endosomal membranes by LLOMe. However, other subunits of ESCRT-0, ESCRT-I or ESCRT-III (CHMP3) were also recruited to varying extents by LLOMe (Fig 2m-n vs Fig 1l and Fig S2c, S3). Much like after hypertonic shock, Lyso Flipper lifetime decreased after LLOMe treatment, indicating that endosomal membrane tension decreased (Fig 2o, Fig S5c). By contrast, the membrane tension of RAB5Q79L early endosomes was not affected by LLOMe (Fig 2p, Fig S5b), consistent with the fact that LLOMe selectively targets late and acidic endosomal compartments ^16^.

Depletion of TSG101 by RNAi prevented both membrane recruitment of CHMP4B (Fig 2q, Fig S4, Fig S6f) and re-acidification (Fig 2j, Fig S6e) in LLOMe-treated cells, confirming that ESCRT-III membrane recruitment is required to repair membrane damage. Altogether, these data indicate that ESCRT-III is recruited onto late endocytic compartments after membrane damage, presumably because membrane tension was relaxed. However, it is also possible that the membrane pores generated by LLOMe directly recruit ESCRT-III rather than decreased tension. We thus searched for a direct approach to show that a decrease in membrane tension promotes assembly of ESCRT-III molecules to the membrane.

To this end, we used a simplified system that by-passes the need for some regulatory factors and is amenable to direct bio-physical manipulations and measurements, consisting of purified, recombinant human CHMP4B labelled with Alexa 488 and giant unilamellar vesicles (GUVs), as model membranes. The GUV lipid composition was dioleoyl-phosphatidylcholine (DOPC):dioleoyl-phosphatidylserine (DOPS) [60:40 Mol%], negatively charged lipid DOPS being required for CHMP4B association to the bilayer ^18^. Upon incubation with 1µM CHMP4B under isotonic conditions (250mOsm), CHMP4B was detected on the bilayer after 30min, consistent with the relatively slow kinetics of nucleation and filament growth ^18^ (Fig 3a-c). Replacement of isotonic with hypertonic solution (500mOsm) caused a three-fold increase in the binding rate of CHMP4B to the GUV (Fig 3a-e, Movie S5) — binding rates can be extracted from exponential fits (Fig 3e). By contrast, almost no CHMP4B binding was observed under hypotonic conditions (Fig 3f). Moreover, following the hypertonic treatment, a hypotonic solution significantly reduced CHMP4B binding (Fig 3c, and see kymograph in Fig 3b), further demonstrating the role of osmotic pressure in the recruitment process. Similar results were obtained with Snf7, the yeast CHMP4 homologue (Fig S7h-i). These data show that hyperosmotic conditions stimulate CHMP4B binding to artificial membranes in vitro, much like on endosomes in vivo.

**Figure 3:**
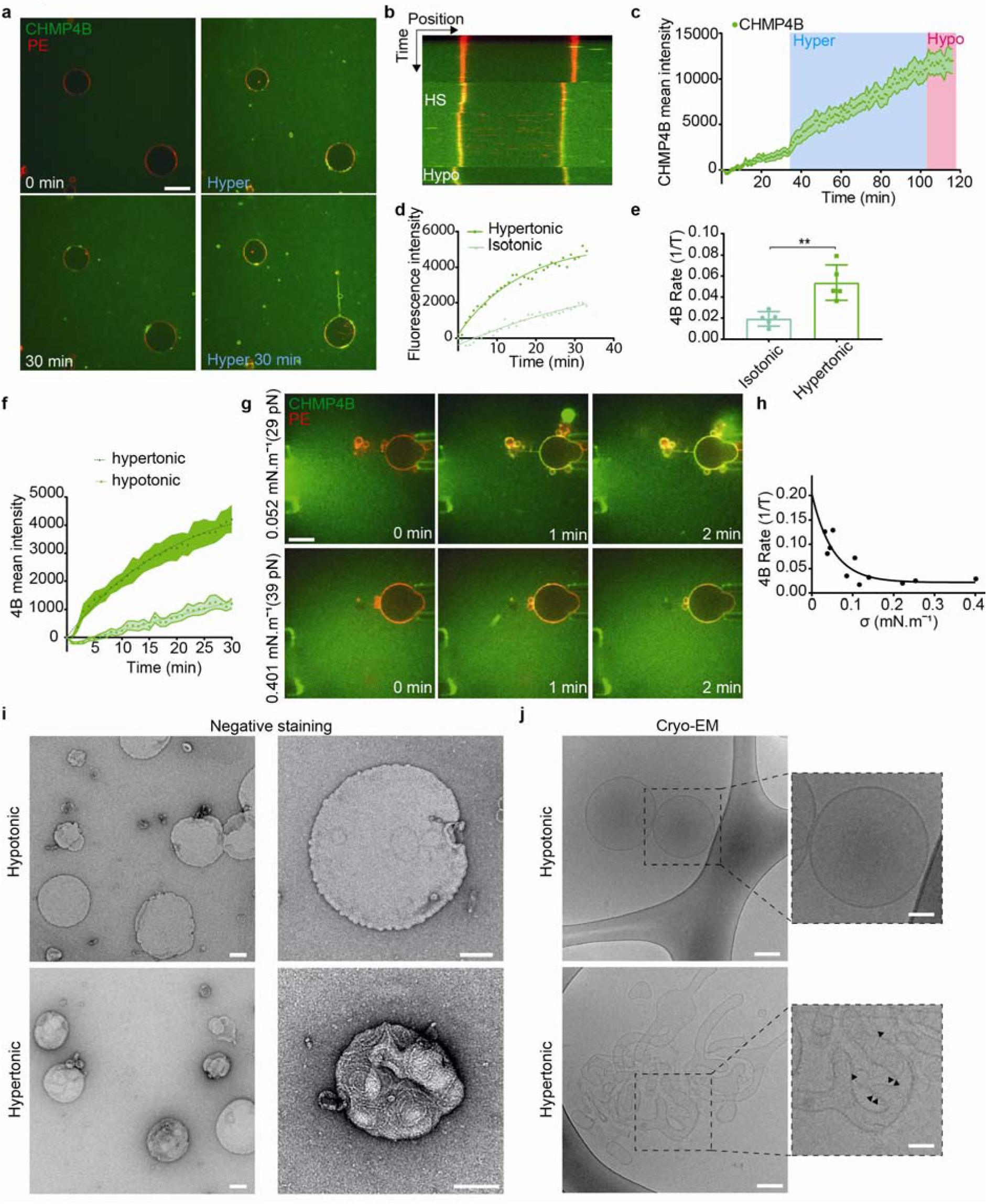
A decrease in membrane tension increases CHMP4B polymerisation rate in vitro. a) Time-lapse confocal images rhodamine-PE labelled GUVs (red) were mixed (0min) with 1 µM CHMP4B-Alexa488 (green), and incubated first in isotonic (250mOsm, 0-30min, white captions) and then in hypertonic (500 mOsm, 0-30min, blue captions) solutions. Bar: 20 µm. b-d) Following experiment shown in (a), GUVs were switched to a hypotonic solution (200 mOsm) for 10min. b) shows a kymograph, c) mean intensity of CHMP4B on the bilayer over time; shaded area: SEM (N=13 GUVs), and d) CHMP4B-Alexa488 mean intensity with time on the GUV during isotonic and hypertonic phases. e) Binding rates extracted from single exponential fits to data as shown in d) (N=6). Each point represents a single experiment, for which 3-13 GUVs were analysed. (two-tailed paired t-test: P=0.0021). f) CHMP4B-Alexa488 mean intensity on GUVs under hypotonic (N=10 GUVs) and hypertonic conditions (N=8 GUVs). Binding rates are: hypotonic, 1/τ=0.0053 (±0.009150) s^-1^; hypertonic, 1/τ=0.05067 (±0.004313) s^-1^. g) The membrane tension of Rhod-PE GUVs (red) was adjusted using a micropipette, and measured by pulling a membrane tube with optical tweezers, while an isotonic solution containing CHMP4B (green) was injected (see diagram in Fig S6a). Time-lapse confocal images show CHMP4B membrane association at low (upper panel), and high membrane tension (lower panel). H) CHMP4B binding rates (1/τ) plotted against membrane tension, as obtained from several experiments (one per dot) as in g). A single exponential decay (black curve) was fitted (R^2^=0.76). I) Negative stain electron micrographs of LUVs incubated with 1µM CHMP4B for 2h in a hypotonic (upper panels) or hypertonic solution (lower panels) at low (left panels) and high (right panels) magnification (Bars: 100 nm). j) Cryo-electron micrographs of LUVs in hypotonic and hypertonic conditions: CHMP4B filaments can be observed under hypertonic conditions (black arrowheads). Bars: left panels, 50 nm; right panels: 25 nm.

To change membrane tension without changing osmolarity, we controlled tension by aspirating GUVs with micropipettes and monitored CHMP4B membrane binding rates (see outline in Fig S7a), while an isotonic solution containing CHMP4B was injected. Then, the aspiration pressure was decreased (Fig S7a), as evidenced by the disappearance of the membrane tongue in the pipette (Fig S7f). The decrease in membrane tension nicely correlated with an almost two-fold increase in CHMP4B binding rate (Fig.S7g). Finally, the dependence of CHMP4B binding rate on tension was quantified directly. Membrane tension was measured by pulling membrane tubes from GUVs ^18^ using optical tweezers ^19^ (Fig 3g). CHMP4B binding rate inversely correlated with membrane tension. An exponential-fit revealed that above a threshold tension of ∼0.1 mN.m^-1^, CHMP4B binding rate was severely reduced (Fig 3h). Interestingly, this threshold value is similar to Chmp4B/Snf7 polymerization energy ^18^, suggesting that tension could directly compete with CHMP4B polymerization ^20^.

To visualize CHMP4B oligomers onto the membrane under varying osmotic conditions, large unilamellar vesicles (LUVs) were incubated with 1µM CHMP4B, negatively stained and analysed by electron microscopy. Under hypertonic conditions, CHMP4B formed spirals (Fig 3i; Fig S8d), resembling the spirals formed by Snf7 in vitro ^18^, and observed with CHMP4B in vivo ^21^ (Fig 3i; Fig S8d). Such spirals were rarely observed under isotonic conditions and almost never under hypotonic conditions (Fig 3i; Fig S8c). These data further confirmed that CHMP4B membrane polymerization is facilitated by a decrease in membrane tension. We hypothesized that membrane deformation coupled to ESCRT-III polymerization is energetically more favourable when membrane tension is low ^18^. In this case, one would expect CHMP4B polymerization to cause membrane deformation.

To test this possibility, CHMP4B-decorated LUVs were analysed by cryo-electron microscopy. In these samples, regularly spaced filaments were clearly visible on the bilayer after incubation under hypertonic conditions, but almost never observed under hypotonic conditions (Fig 3j). Importantly, tubular and vesicular deformations, covered with CHMP4B filaments could be observed in hypertonic solutions, while essentially absent under hypotonic conditions (Fig 3j, Fig S8a-b). Furthermore, LUVs that were not decorated by filaments – even in hypertonic conditions – were not deformed (Fig 3j, Fig S6a-b). This strongly support the view that a decrease in membrane tension facilitates membrane deformation by ESCRT-III polymerization.

Given the role of ESCRT-III in ILV formation, we wondered if a decrease of endosomal membrane tension could trigger ILV formation. First, we analysed the ultrastructure of endosomes after hypertonic shock by Correlative Light-Focused Ion Beam-Scanning Electron Microscopy (FIB-CLEM). CHMP4B-labeled endosomes (Fig 4a) appeared more electron-dense after hypertonic shock when compared to untreated controls. Moreover, the density of endosomes containing BSA-gold endocytosed for 15min (early endosomes) was increased after hypertonic shock when compared to controls (Fig S9a). Late endosomes containing endocytosed BSA gold chased for 2h were more electron-dense than early endosomes even without treatment, and yet the treatment also increased their density (Fig S9a). While this increase may be due to volume reduction (Fig 2a-d), we reasoned that it may also result from an increased number of ILVs formed upon hypertonic shock.

**Figure 4:**
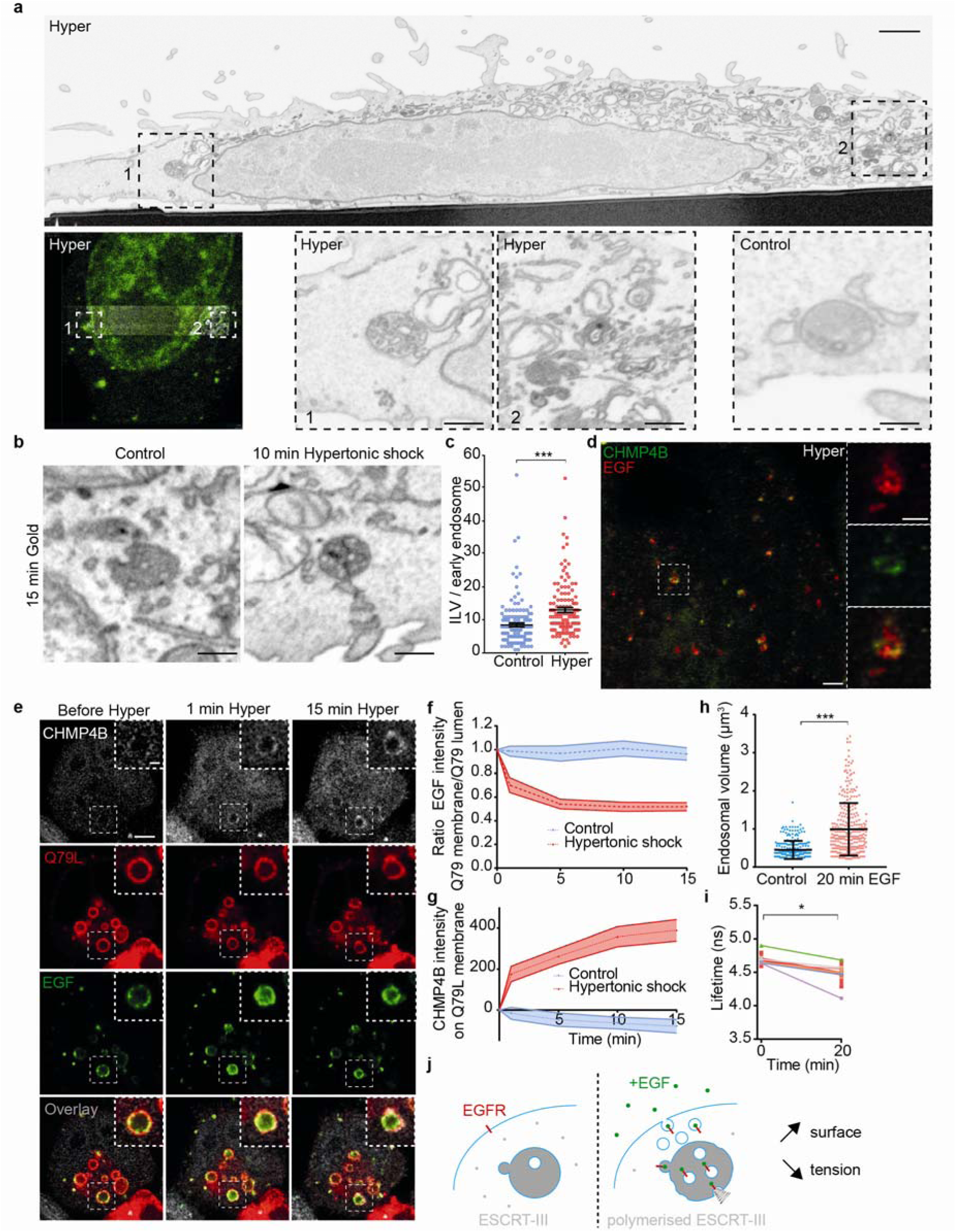
Tension-induced CHMP4B recruitment on endosomes triggers ILV formation. a) CLEM-FIB-SEM micrographs of cells expressing CHMP4B-GFP treated with hypertonic medium (800 mOsm) for 10min. A whole cell section (upper panel, bar: 5 µm) shows CHMP4B-GFP decorated endosomes (left panel) in boxed areas 1 and 2 (high magnification in lower panels, bar 1 µm). Lower control panel shows a typical endosome under isotonic condition (bar: 200 nm). b-c) From FIB-SEM micrographs (b) of cells loaded with BSA-gold, ILV number per BSA-gold labelled early endosomes (EE) was quantified before (blue) and after (red) a 10min hypertonic shock Bars: 200 nm (c). Error bar is SEM (Control: 152, Hyper: 131 endosomes in 3 independent replicates, two-tailed Mann-Whitney test: P=0.0000000002). d) STED microscopy images of cells expressing CHMP4B-GFP (green), incubated 10 minutes with far red EGF (red) and then subjected to 10min hypertonic shock. Bar: 2 μm High magnification (right panels) bars: 500 nm. e) Time-lapse confocal images of Hela-CHMP4B-GFP cells overexpressing RAB5Q79L-mcherry incubated 10 minutes with EGF-Alexa647, and subjected to hypertonic shock. f) Ratio of EGF fluorescence colocalizing with RAB5Q79L membrane over total EGF signal, normalized by the initial value. g) Intensity of CHMP4B-GFP on Rab5Q79L membrane. For g) and f), shaded areas are SEM (N=21 endosomes from 3 independent replicates (hypertonic shock) and N=16 endosomes from 3 independent replicates (control condition, isotonic). h) Volumes of endosomes stained with Lyso Flipper before and 20min after 200 ng/ml EGF treatment. Dots correspond to single endosomes. Black line: Mean±SD (N=266 endosomes (before) and N=308 (Hyper) from 3 independent replicates. Two-tailed Mann-Whitney: P<10^-15^). i) Lyso Flipper lifetime measurements before and 20min after 200 ng/ml EGF treatment. Thin lines: 5 independent replicates (a few hundreds of endosomes from more than 3 cells each); thick red line: mean±SEM, two-tailed paired t-test P=0.025138. j) Schematic of a putative mechanism for membrane tension dependent ILV formation by ESCRT-III machinery (see text).

In FIB-SEM samples, the number of ILVs indeed increased by almost 40% in early endosomes after hypertonic shock (Fig 4b,c), supporting the view that ESCRT-III recruitment under hypertonic conditions triggers ILV formation. Furthermore, CHMP4B co-clustered with EGF on endosomes (Fig 4d) – presumably ILV buds – as seen by STED. To test whether ILVs formed upon hypertonic shock, we monitored EGF receptor sorting into enlarged RAB5Q79L endosomes, which provide sufficient space resolution ^12,22^. Cells were incubated with fluorescent EGF for 10min, and then EGF fluorescence intensity on the limiting membrane of enlarged endosomes was measured before and after hypertonic shock. Upon shock, the fluorescence intensity of CHMP4B increased (Fig 4e,g), as expected (Fig 2a), concomitant with an increase of the EGF signal in the endosomal lumen (Fig 4e). This increase presumably reflects EGFR sorting into ILVs, although it may also reflect an increased concentration due to the decreased endosome volume (Fig 2b-d). We thus monitored the decrease in EGF signal on the limiting membrane (Fig 4f), since this can only be accounted by ILV formation — membrane traffic being inhibited under hypertonic conditions. Altogether, these observations show that a decrease in membrane tension causes the recruitment of CHMP4B on endosomal membranes, which in turn drives the formation of intralumenal vesicles containing the EGF receptor.

These results suggested that membrane tension regulation may be at play during physiological trafficking of the EGF receptor. Indeed, EGF induces massive endocytosis of EGF receptor and stimulates ESCRT-dependent ILV formation in endosomes ^23^. Therefore, we measured membrane tension of endosomes using Lyso Flipper after EGF treatment. Interestingly, 20min after 200ng/ml EGF addition, the endosomal volume increased almost two fold (Fig 4h), as previously shown ^24^. More importantly, endosomal membrane tension was significantly reduced (Fig 4i, Fig S9b) showing that it is negatively regulated upon EGF-dependent ILV formation under physiological conditions. We propose that the fusion of endocytic vesicles containing the EGF receptor with endosomes increases the endosome membrane surface area, which in turn decreases membrane tension and promotes ILV formation by the ESCRT machinery (Fig 4j).

In conclusion, our in vitro data unambiguously demonstrate that CHMP4B can polymerize on the membrane only when membrane tension is lowered. The precise role of lipid organization and packing in CHMP4B membrane association remains unclear. However, it is tempting to speculate that decreased tension, which influences lipid packing and lateral mobility ^25^, directly facilitates membrane interactions with CHMP4B. Clearly, it can be expected that in vivo such interactions are stabilized by the coincident detection of multiple binding partners ^26^ and protein-protein interactions. Indeed, we find that both ALIX and TSG101, which interact physically with each other ^27^, act as ESCRT-III nucleator in vivo. Our data indicate that CHMP4B recruitment under hypertonic conditions and LLOMe treatment exhibit some preference for ALIX and TSG101, respectively, perhaps reflecting subtle changes in membrane organization — hence in the capacity to interact with ALIX or TSG101 — after either treatment.

As CHMP4B uses membranes as a substrate to polymerize, a lower tension likely facilitates ESCRT polymerisation in its preferred curvature radius. Conversely, high tension may inhibit CHMP4B binding as the polymerisation energy will not be sufficient to allow membrane deformation. It is tempting to speculate that this mechanism is shared by all ESCRT-III mediated reactions, and that membrane tension drives all ESCRT-III dependent processes.

## ACKNOWLEDGEMENTS

The authors want to thank ACCESS, the Bioimaging platforms, bioimaging core facility and the electron microscopy core facility of University of Geneva for constant support. JG acknowledges support from the Swiss National Science Foundation Grant No 31003A_159479, the NCCR in Chemical Biology and LipidX from the Swiss SystemsX.ch Initiative. AR acknowledges funding from Human Frontier Science Program Young Investigator Grant RGY0076/2009-C, the Swiss National Fund for Research Grants N°31003A_130520, N°31003A_149975 and N°31003A_173087, and the European Research Council Consolidator Grant N° 311536.

## DATA AVAILABILITY

The data that support the findings of this study are available from the corresponding authors upon reasonable request.

## AUTHOR CONTRIBUTIONS

VM, JG and AR designed the project based on the first observation made by GM. VM carried out all experiments and analyses. JL performed some experiments with VM, in particular CHMP4B purification. AG and SM designed and synthesised the Lyso Flipper and FliptR molecules. VM, JG and AR wrote the paper, with corrections from all co-authors.

## References and Notes

1 Pontes, B., Monzo, P. & Gauthier, N. C. Membrane tension: A challenging but universal physical parameter in cell biology. Seminars in Cell & Developmental Biology 71, 30–41, doi:https://doi.org/10.1016/j.semcdb.2017.08.030 (2017).

2 Vietri, M. et al. Spastin and ESCRT-III coordinate mitotic spindle disassembly and nuclear envelope sealing. Nature 522, 231–235, doi:10.1038/nature14408 (2015).

3 Olmos, Y., Hodgson, L., Mantell, J., Verkade, P. & Carlton, J. G. ESCRT-III controls nuclear envelope reformation. Nature 522, 236–239, doi:10.1038/nature14503 (2015).

4 Denais, C. M. et al. Nuclear envelope rupture and repair during cancer cell migration. Science 352, 353 (2016).

5 Raab, M. et al. ESCRT III repairs nuclear envelope ruptures during cell migration to limit DNA damage and cell death. Science 352, 359 (2016).

6 Jimenez, A. J. et al. ESCRT machinery is required for plasma membrane repair. Science 343, 1247136, doi:10.1126/science.1247136 (2014).

7 Skowyra, M. L., Schlesinger, P. H., Naismith, T. V. & Hanson, P. I. Triggered recruitment of ESCRT machinery promotes endolysosomal repair. Science 360 (2018).

8 Colom, A. et al. A fluorescent membrane tension probe. Nature Chemistry 10, 1118–1125, doi:10.1038/s41557-018-0127-3 (2018).

9 Mierzwa, B. E. et al. Dynamic subunit turnover in ESCRT-III assemblies is regulated by Vps4 to mediate membrane remodelling during cytokinesis. Nat Cell Biol 19, 787–798, doi:10.1038/ncb3559 (2017).

10 Bissig, C. et al. Viral infection controlled by a calcium-dependent lipid-binding module in ALIX. Dev Cell 25, 364–373, doi:10.1016/j.devcel.2013.04.003 (2013).

11 Raiborg, C. et al. Hrs sorts ubiquitinated proteins into clathrin-coated microdomains of early endosomes. Nat Cell Biol 4, 394–398, doi:10.1038/ncb791 (2002).

12 Pons, V. et al. Hrs and SNX3 Functions in Sorting and Membrane Invagination within Multivesicular Bodies. PLOS Biology 6, e214, doi:10.1371/journal.pbio.0060214 (2008).

13 Heuser, J. E. & Anderson, R. G. Hypertonic media inhibit receptor-mediated endocytosis by blocking clathrin-coated pit formation. The Journal of cell biology 108, 389–400, doi:10.1083/jcb.108.2.389 (1989).

14 Nunes, P. et al. Ionic imbalance, in addition to molecular crowding, abates cytoskeletal dynamics and vesicle motility during hypertonic stress. Proceedings of the National Academy of Sciences 112, E3104, doi:10.1073/pnas.1421290112 (2015).

15 Goujon, A. et al. Mechanosensitive Fluorescent Probes to Image Membrane Tension in Mitochondria, Endoplasmic Reticulum, and Lysosomes. Journal of the American Chemical Society, doi:10.1021/jacs.8b13189 (2019).

16 Repnik, U. et al. LLOMe does not release cysteine cathepsins to the cytosol but inactivates them in transiently permeabilized lysosomes. Journal of Cell Science (2017).

17 Radulovic, M. et al. ESCRT-mediated lysosome repair precedes lysophagy and promotes cell survival. The EMBO Journal 37, doi:10.15252/embj.201899753 (2018).

18 Chiaruttini, N. et al. Relaxation of Loaded ESCRT-III Spiral Springs Drives Membrane Deformation. Cell 163, 866–879, doi:10.1016/j.cell.2015.10.017 (2015).

19 Morlot, S. et al. Membrane shape at the edge of the dynamin helix sets location and duration of the fission reaction. Cell 151, 619–629, doi:10.1016/j.cell.2012.09.017 (2012).

20 Lenz, M., Crow, D. J. & Joanny, J. F. Membrane buckling induced by curved filaments. Phys Rev Lett 103, 038101, doi:10.1103/PhysRevLett.103.038101 (2009).

21 Hanson, P. I., Roth, R., Lin, Y. & Heuser, J. E. Plasma membrane deformation by circular arrays of ESCRT-III protein filaments. J Cell Biol 180, 389–402, doi:10.1083/jcb.200707031 (2008).

22 Bache, K. G. et al. The ESCRT-III subunit hVps24 is required for degradation but not silencing of the epidermal growth factor receptor. Mol Biol Cell 17, 2513–2523, doi:10.1091/mbc.E05-10-0915 (2006).

23 Wenzel, E. M. et al. Concerted ESCRT and clathrin recruitment waves define the timing and morphology of intraluminal vesicle formation. Nature communications 9, 2932–2932, doi:10.1038/s41467-018-05345-8 (2018).

24 Razi, M. & Futter, C. E. Distinct roles for Tsg101 and Hrs in multivesicular body formation and inward vesiculation. Mol Biol Cell 17, 3469–3483, doi:10.1091/mbc.E05-11-1054 (2006).

25 Barelli, H. & Antonny, B. Lipid unsaturation and organelle dynamics. Current Opinion in Cell Biology 41, 25–32, doi:https://doi.org/10.1016/j.ceb.2016.03.012 (2016).

26 Carlton, J. G. & Cullen, P. J. Coincidence detection in phosphoinositide signaling. Trends in cell biology 15, 540–547, doi:10.1016/j.tcb.2005.08.005 (2005).

27 von Schwedler, U. K. et al. The Protein Network of HIV Budding. Cell 114, 701–713, doi:http://dx.doi.org/10.1016/S0092-8674(03)00714-1 (2003).

