## Supplementary material for "Endosomal Membrane Tension Regulates ESCRT-III-Dependent Intra-Lumenal Vesicle Formation"

*co-corresponding and co-last authors

**This PDF file includes:**

Materials and Methods

Figs. S1 to S9

Captions for Movies S1 to S5

**Other Supplementary Materials for this manuscript include the following:**

Movies S1 to S5

Materials and Methods

**Cells, Antibodies, Reagents**

HeLa-MZ cells (Lucas Pelkmans, University of Zurich), HeLa Kyoto cells stably expressing GFP-CHMP4B-GFP (Anthony Hyman, MPI‐CBG, Dresden), MDA MB 431 cells were maintained as described (*1*).HeLa and MDA MB 431 cells are not on the list of commonly misidentified cell lines maintained by the International Cell Line Authentication Committee. Our HeLa-MZ, HeLa Kyoto and MDA MB 431 cells were authenticated by Microsynth (Balgach, Switzerland), and are mycoplasma-negative as tested by GATC Biotech (Konstanz, Germany).

Mouse monoclonal antibodies against LBPA (*2*). and Rab5 (*3*) were previously described. Rabbit monoclonal antibodies against Lamp1 (D2D11, 9091) were from Cell Signaling (Danvers, MA USA), the rabbit polyclonal antibody against EEA1 (ALX-210-239) from ENZO life science (New-York, USA), the mouse monoclonal antibody against EEA1 from BD biosciences (Franklin Lakes, NJ USA), and the anti-tubulin antibody (T9026) from Sigma-Aldrich (St. Louis, MO). Antibodies against CHMP1A (15761-1-AP), CHMP2A (10477-1-AP), CHMP1B (14639-1-AP), CHMP2B (12527-1-AP), CHMP4B (13683-1-AP), Alix (12422-1-AP), VPS4B (17673-1-AP), VPS25 (15669-1-AP), STAM1 (12434-1-AP), TSG101 (14497-1-AP), CHMP6 (16278-1-AP) and CHMP3 (15472-1-AP) were from Proteintech (Rosemont, IL, USA).

We obtained FM4-64, LysoTracker, Alexa 647-EGF, Alexa-555 EGF and TRITC-dextran (T13320, L7528, E35351, E35350 and D1817, respectively) from Thermo Fisher Scientific (Waltham, MA), as well as EZ-Link Sulfo-NHS-SS-Biotin, Alexa Fluor 488 TFP ester, Ni-NTA Agarose, 7 kDa MWCO Zeba Spin Desalting Columns and Hoechst 33342. Peroxidase-conjugated secondary antibodies were from Bio-Rad Laboratories (Hercules, CA), MBPTrap HD 5mL columns, Glutathione Sepharose 4B and Dextrin Sepharose High Performance from GE Healthcare (Anaheim, CA), and the cOmplete Protease Inhibitor Cocktail from Roche (Basel, Switzerland). We obtained DOPC, DSPE-PEG-Biotin, DOPE, DOPS and lissamine rhodamine B sulfonyl (18:1 Rhod PE from Avanti Polar Lipids (Alabaster, AL, and the lysosomotropic reagent L-leucyl-L-leucine methyl ester (LLOMe) from Sigma-Aldrich (St. Louis, MO). Other reagents and chemicals were obtained from Sigma-Aldrich (St. Louis, MO).

**siRNA treatment**

siRNA sequences against Alix and TSG101 were from siTOOLs Biotech (Planegg, Germany). For each gene, a pool of 30 different siRNA sequences was designed. A siRNA sequence against VSV-G, which does not interact with any human genes, was used as a negative control. Each pool was used at a low 3nM (20 nM for anti-VSV-G siRNA) concentration in order to reduce the danger of off-target effects. DNA and siRNA were transfected in cells according to the manufacturer’s instructions using FuGENE HD (Promega Corporation, Madison, WI) and Lipofectamine RNAiMAX (Thermo Fisher Scientific, Waltham, MA), respectively. Unless indicated otherwise, experiments were performed after transfection with DNA for 7h and siRNA for 72 h.

**Live-cell imaging and Immunofluorescence**

For live-cell imaging, cells were seeded into 35mm MatTek glass bottom microwell dishes (MatTek Corporation) or in Ibidi multi-wells chambers (Ibidi) with 24x60mm glass coverslips (Menzel-Glaser). For immunofluorescence studies, cells were seeded at 4.10^4^ cells per 35 mm dish and grown on 1.5 round glass coverslips (Electron Microscopy Sciences, Hatfield, PA, USA) at 37°C or 5.10^3^ cells per well in Ibidi p96 wells plate (or 2. 10^3^ cells per well for siRNA experiment). For live imaging, cells were rinsed 3 fold with 1ml Leibovitz’s live imaging medium (Life Technologies, Thermofisher so that cells could be incubated without CO_2_ equilibration at room temperature. Long (>30 min) acquisitions were at 37°C in Leibovitz’s medium. For EEA1, LBPA and Lamp1, immunofluorescence staining was performed as described (*4*).

For ESCRT staining, cells were fixed with methanol during 10min at -20°C and blocked using 1% p/v BSA in PBS. ESCRT were then immunolabeled in this blocking solution for 1h at room temperature with antibodies against the following proteins:, CHMP1B, CHMP3, CHMP4B, Alix, VPS4B, TSG101, STAM (All were rabbit polyclonal antibodies obtained from ProteinTech Group, Rosemont, IL, USA, with the following reference numbers, respectively: 14639-1-AP, 15472-1-AP, 13683-1-AP, 12422-1-AP, 17673-1-AP, 14497-1-AP, 12434-1-AP) all at a 300-fold dilution. Otherwise, the labelling protocol was as described (*4*).

Cells were analysed by confocal microscopy (Leica SP8, Nikon A1), STED microscopy (Leica SP 8) or confocal automated microscope (Molecular Device) for P96 wells plates. Analysis of automated microscopy images was done via the MetaXpress Custom Module editor software Images were segmented in order to generate relevant masks, which were then applied on the fluorescent images to extract relevant measurements. To facilitate segmentation of ESCRT or LysoTracker endosomes, we applied the top hat deconvolution method, when necessary, to reduce the background noise and highlight bright granules.

**Giant unilamellar vesicles (GUVs) and large unilamellar vesicles (LUVs) preparation**

GUVs were produced as described (*5*). Here, the lipid composition was DOPC:DOPS at 60:40 Mol%, containing 0.1% N-rhodamine PE and 0.003% DSPE-PEG(2000) Biotin. To prepare LUVs, 500 µl of the same lipid mix than for GUVs (2mg/ml) were evaporated in round bottom glass tubes under gentle CO_2_ flow. Then, 500 µl water containing 250mM sucrose were added onto the lipid film and the mixture was vortexed during 5min. The solution was size-extruded using 400nM filter to form LUVs.

**CHMP4B binding on GUVs**

A channel of ibidi flow chamber was coated with 0.1 mg/ml avidin in water for 10min, and rinsed 3X with water and 2X with buffer 1 (100mM NaCl, 20mM HEPES, 2mM MgCl2, 250mOsm). 2µl GUVs were added in 50µl buffer 1 during 5min to facilitate avidin-biotin interactions, and then the solution was replaced by 40µl of 0.1mg/ml albumin-biotin in buffer 1 in order to saturate free avidin sites. For analysis, ROIs were selected by focusing on the GUV equatorial plane at 1 frame/1-2min, and acquisition started before CHMP4B-Alexa 488 addition. In the experiment, CHMP4B-Alexa 488 was diluted 1:1 with buffer 2 (200mM NaCl, 20mM HEPES, 4mM MgCl_2_). The mixture was adjusted to 1µM CHMP4B with buffer 1 (CHMP4B Iso), and 80µl were added to GUVs with a pump at a rate = 20µl/min to avoid GUV damages. For hyperosmotic CHMP4B incubation, protein was mixed with buffer 1 and 2 supplemented with 250mM sucrose (CHMP4B Hyper, final osmolarity ~500mOsm). For hypotonic shock, the hypertonic solution was replaced by 1µM CHMP4B in buffer 1 supplemented with water to reach ~230mOsm (instead of ≈250mOsm for isotonic solution). For quantification, the red channel (membrane) was threshold and used as a mask for the green (CHMP4B) signal over time. Background was subtracted and curves were fitted using one phase association exponential (formula: *y* = *y0+(plateau-y0)(1-* *e*^−^*^Kx^*)) in order to extract binding rates.

**Tube pulling experiments**

The experimental setup used to aspirate GUV with a micropipette and pull a membrane tube was the same than (*5*), and is combining brightfield imaging, spinning disc confocal microscopy and optical tweezers on an inverted Nikon Eclipse Ti microscope. The force F exerted on the bead by the membrane tube was calculated from Hooke’s law: F = k.Δx, where k is the stiffness of the trap (k = 60 pN.μm^−1^) and Δx the displacement of the bead from its equilibrium position in the optical trap. CHMP4B-A488 was injected with a micropipette connected to a motorized micromanipulator and to the Fluigent pressure control system (MFCS-VAC −69 mbar, Fluigent, Villejuif, France).

Binding rates of CHMP4B on GUVs membrane were extracted using one phase association exponential (formula: *y* = *y0+(plateau-y0)(1-* *e*^−^*^Kx^*)). Finally, we plotted the binding rates of CHMP4B as a function of membrane tension (extracted from equation: F=2π√2κσ) and we fitted this curve with one phase exponential decay (*y=(y0 – plateau) ).e^−Kx^ + Plateau*; R²=0.77)

**Cryo-electron microscopy and negative-staining transmission electron microscopy**

In cryo-electron microscopy studies, LUVs were adsorbed for 1min onto EM grids with 300 mesh Lacey carbon films, blotted with Whatman filter paper, and then plunged in liquid ethane using FEI vitrobot cryoplunger (FEI, Eindhoven, the Netherlands). Cryo-electron micrographs were recorded using a field emission gun transmission electron microscope Tecnai F 20 (FEI, Eindhoven, the Netherlands) equipped with a cryo-specimen grid holder Gatan 626 (Warrendale, Pennsylvania, USA). Images were acquired at 200kV and at a nominal magnification of 50,000 using an Eagle camera (4,096 pixels×4,096 pixels). Samples were analysed by negative-staining transmission electron microscopy, as described (*5*).

**Endosomes purification**

Subcellular fractionation was carried out by flotation in sucrose gradients as described (*6*). Whole cell lysates were prepared in 50 mM Tris, pH 7.4, 1% NP-40, 0.25% sodium deoxycholate, 150 mM NaCl, 1mM EDTA, 1mM PMSF, 1 μg/mL aprotinin, 1 μg/mL leupeptin, 1 μg/mL pepstatin.

**Fluorescence Lifetime Imaging Microscopy (FLIM)**

FliptR and Lyso Flipper were synthesized following reported procedures (*7, 8*) and FLIM with these probes was adapted from (*7, 8*). Cells were incubated with 1μM FliptR (for RAB5Q79L endosome staining) during 2h before imaging or with 1μM Lyso FlipperLyso Flipper during 15min before imaging. Imaging was performed as in (*8*). For analysis, SymPhoTime 64 software (PicoQuant) was used to fit fluorescence decay data (from full images for Lyso FlipperLyso Flipper or regions of interest for FliptR) to a dual exponential reconvolution model whereby the lifetime τ1 was extracted. Data are expressed as means ± SD.

**FRAP experiment**

Endosomes were imaged using a 100x 1.4 NA oil DIC Plan-Apochromat VC objective (Nikon) with a Nikon A1 scanning confocal microscope. Photobleaching were performed in circular regions with 4 iterations of 488nm at 100% laser intensity. For the analysis, the fluorescence background was subtracted from ROIs intensity values. Those values were subsequently normalized by a non-bleached area and the value of intensity of the first frame after bleaching was subtracted. Finally, FRAP curves were normalized with the pre-bleach fluorescence intensity value. To determine FRAP recovery kinetics, we fitted with the following double exponential function  *f(t)=α(1-e^t-t0/τ1^)+(1-α)(1-e^t-t0/τ2^).*

**STED nanoscopy**

Hela-CHMP4B-GFP cells were growth on 1.5cm 170μm thick glass coverslip (Hecht-Assistant), incubated 10min with EGF-A647 in medium at 37°C then rinsed and further incubated for 10min in medium containing 0.5M sucrose at 37°C. Cells were then fixed for 15min at room temperature with 3%PFA in PBS, rinsed 3X with PBS then 1X with water and mounted using Prolong gold mounting medium (ThermoFisher) and sealed using nail polish (Electron Microscopy Sciences).

Image acquisition was performed using SP8 confocal microscope (Leica) equipped with 2 depletion laser lines one at 592nm and the other at 775nm and 93X glycerol PL Apo objective with 1.3 NA. Prior to image the samples, depletion lasers were aligned using gold beads. Then, imaging CHMP4-GFP signal was depleted using 40% power of the 592nm line and EGF-A647 was depleted using 35% of 775 nm laser line. No gating was used for both of them. Images were post processed automatically via the lightning mode of Las X software (Leica) which basically consist of a deconvolution process using the estimated PSF depending on imaging parameters.

**Endosomal volume measurements**

Z-sections were acquired on living cells before and after hypertonic shock using 100X objective with 1.4 numerical aperture on Nikon Ti scanning confocal inverted microscope or Nikon Ti spinning disk inverted microscope. Images were processed with Icy software where endosomes were manually threshold and exported as ROIs. Those ROIs were then convexified and the volumes of ROIs were extracted using the 3D measurment plugin.

**Fluorescent EGF in RabQ79L membrane measurements**

The analysis of EGF receptor distribution on endosomes enlarged after expression of RAB5Q79L, EGF-A647 was endocytosed 10min at 37°C in Hela CHMP4B-GFP expressing RAB5Q79L cherry and z slices were acquired using Nikon A1 microscope before and at indicated times of hypertonic shock. For analysis, EGF signal was measured using Icy software on the limiting membrane of enlarged endosomes and normalized with the cytosolic signal of CHMP4B-GFP and finally expressed as a ratio of signal before hypertonic shock. The CHMP4B-GFP signal was also measured using the same procedure.

**FIB-SEM sample preparation and CLEM**

Briefly, for early endosome staining, Hela cells grown in 35mm plastic dishes were incubated with 6nm BSA-gold particles for 10min at 37 °C, washed extensively with medium and then either fixed immediately or treated for 10min at 37°C with hypertonic medium (0.5M sucrose) before fixation. For late endosome staining, BSA-gold was endocytosed as above for 10min, and then cells were washed extensively with medium, and further incubated at 37 °C for 2h. Cells were then fixed immediately or treated for 10min at 37°C with hypertonic medium (0.5 M sucrose). Cells were rinsed once with PBS and fixed for 3h on ice using 2.5% glutaraldehyde, 2% paraformaldehyde in buffer A containing 0.15M cacodylate and 2mM CaCl2. After this step, cells were processed for FIB-SEM as described (*9*). At the end of sample preparation, a small bloc was cut with an electric saw and incubated for ≈30min in 100% xylene in order to remove plastic leftovers. Finally, the ROI was processed using HELIOS 660 Nanolab DualBeam SEM/FIB (FEI Company, Eindhoven, Netherlands). The ROI size was ≈15-20µm wide and the thickness of the FIB slice was either 5 or 10nm depending on the experiment.

For endosomal density quantification, gold containing endosomes were identified. The image intensity inverted and the mean intensity of endosome were measured. Finally, the intensity of the cytoplasm for each slice was subtracted. Results were analysed using GraphPad Prism. The Gaussian distribution of the data were tested using Kolmogorov–Smirnov test (with Dallal–Wilkinson–Lillie for P value).

For CLEM, 5.10^4^ Hela CHMP4B cells were grown on 35mm dishes with plastic coverslips bottom labelled with grids (Ibidi) for 24h. One dish was left untreated and the other was treated with medium containing 0.5M sucrose during 10min at 37°C. Cells were then fixed with 3% PFA and Z-stacks were acquired using Leica SP8 confocal microscope. Cells were then stained and embedded in Epoxy resin for FIB-SEM acquisition as described above. Using the grid, cells imaged by confocal were found back and imaged using Helios nanolab microscope. Both optical and electronic images were processed using Big Warp plugin on ImageJ and Icy software.

**Cryo-sectioning and immuno-electron microscopy**

Cells were incubated 15 min at 37°C with 6 nm BSA-Gold washed and directly fixed (control) or incubated further for 10 minutes at 37°C in sucrose containing hypertonic medium (830 mOsm) before fixation (10 min hyper). Fixation was carried out fo 2h at room temperature in 0.125M phosphate buffer (PB) containing 2% PFA and 0.2% glutaraldehyde. After ﬁxation, cells were washed 3 times with PB and once with PB containing 50 mM glycine, embedded in 10% (wt/vol) gelatin and infused with 2.3 M sucrose. Small pieces of the cell pellet were then cut and incubated overnight in 2.3 M sucrose. Gelatin blocks were frozen in liquid nitrogen and ultrathin cryo sections were cut (35 nm) at −120°C using an Ultracut FCS ultracryomicrotome (Leica-Reichert). Sections were retrieved in a 1:1 solution of 2.3 M sucrose and 2% methyl cellulose. For labeling, cryosections were first blocked in PBS containing 1% BSA and incubated with primary anti-CHMP4B antibody (Protein tech, 13683-1-AP) diluted 30 times in the same solution. Cryosections were then washed 4 times in PBS and incubated with a solution of 10 nm gold-conjugated protein A (AURION) at OD = 1 in 1% BSA PBS. Finally, cryosections were stained with 1% uranyl acetate in 2% methylcellulose. The sections were viewed at 120 kV in a Technai G2 Sphera electron microscope.

**Statistical Analyses**

Results shown are means, and error bars represent standard errors of the mean (SEM) or standard deviation (SD) as specified in figure legends. Except where otherwise stated, experiments were repeated at least 3 times. Statistical analyses were performed using Prism 7 (GraphPad Software). The Gaussian distribution of the data were tested using Kolmogorov–Smirnov test (with Dallal–Wilkinson–Lillie for P value). In case of non-Gaussian distribution, the following non-parametric tests were used: two-tailed Mann–Whitney U test to compare two conditions, or one-way ANOVA (Kruskal–Wallis test) with a Dunn’s test to compare more than two data groups. In case of Gaussian distribution, two-tailed Student’s t-test was used for the comparison of the means in case of comparison between two conditions and parametric one-way ANOVA with a Tukey HSD or Dunnett’s multiple comparison tests in case of comparisons between more than two data groups. When appropriate, correlation coefficients were calculated using either Pearson or Spearman. Significance of mean comparison is represented on the graphs by asterisks: *<0.05; **<0.01; ***<0.001.

**Other methods**

Western blot analysis was preformed as (*4*), and blot exposure times were always within the linear range of detection. Human CHMP4B was purified as described (*10*). The analysis of EGF receptor endocytosis was described (*11*) and endocytosis of BSA-gold was described (*4*).

**Supplementary References**

1.	C. C. Scott, S. Vossio, J. Rougemont, J. Gruenberg, TFAP2 transcription factors are regulators of lipid droplet biogenesis. *eLife* **7**, e36330 (2018).

2.	T. Kobayashi *et al.*, A lipid associated with the antiphospholipid syndrome regulates endosome structure and function. *Nature* **392**, 193-197 (1998).

3.	V. Cavalli *et al.*, The Stress-Induced MAP Kinase p38 Regulates Endocytic Trafficking via the GDI:Rab5 Complex. *Molecular Cell* **7**, 421-432 (2001).

4.	V. Mercier *et al.*, ALG-2 interacting protein-X (Alix) is essential for clathrin-independent endocytosis and signaling. *Sci Rep* **6**, 26986 (2016).

5.	N. Chiaruttini *et al.*, Relaxation of Loaded ESCRT-III Spiral Springs Drives Membrane Deformation. *Cell* **163**, 866-879 (2015).

6.	J.-P. Gorvel, P. Chavrier, M. Zerial, J. Gruenberg, rab5 controls early endosome fusion in vitro. *Cell* **64**, 915-925 (1991).

7.	A. Goujon *et al.*, Mechanosensitive Fluorescent Probes to Image Membrane Tension in Mitochondria, Endoplasmic Reticulum, and Lysosomes. *Journal of the American Chemical Society*, (2019).

8.	A. Colom *et al.*, A fluorescent membrane tension probe. *Nature Chemistry* **10**, 1118-1125 (2018).

9.	P. Nunes-Hasler *et al.*, STIM1 promotes migration, phagosomal maturation and antigen cross-presentation in dendritic cells. *Nature communications* **8**, 1852-1852 (2017).

10.	B. E. Mierzwa *et al.*, Dynamic subunit turnover in ESCRT-III assemblies is regulated by Vps4 to mediate membrane remodelling during cytokinesis. *Nat Cell Biol* **19**, 787-798 (2017).

11.	B. Brankatschk *et al.*, Regulation of the EGF Transcriptional Response by Endocytic Sorting. *Science Signaling* **5**, ra21 (2012).

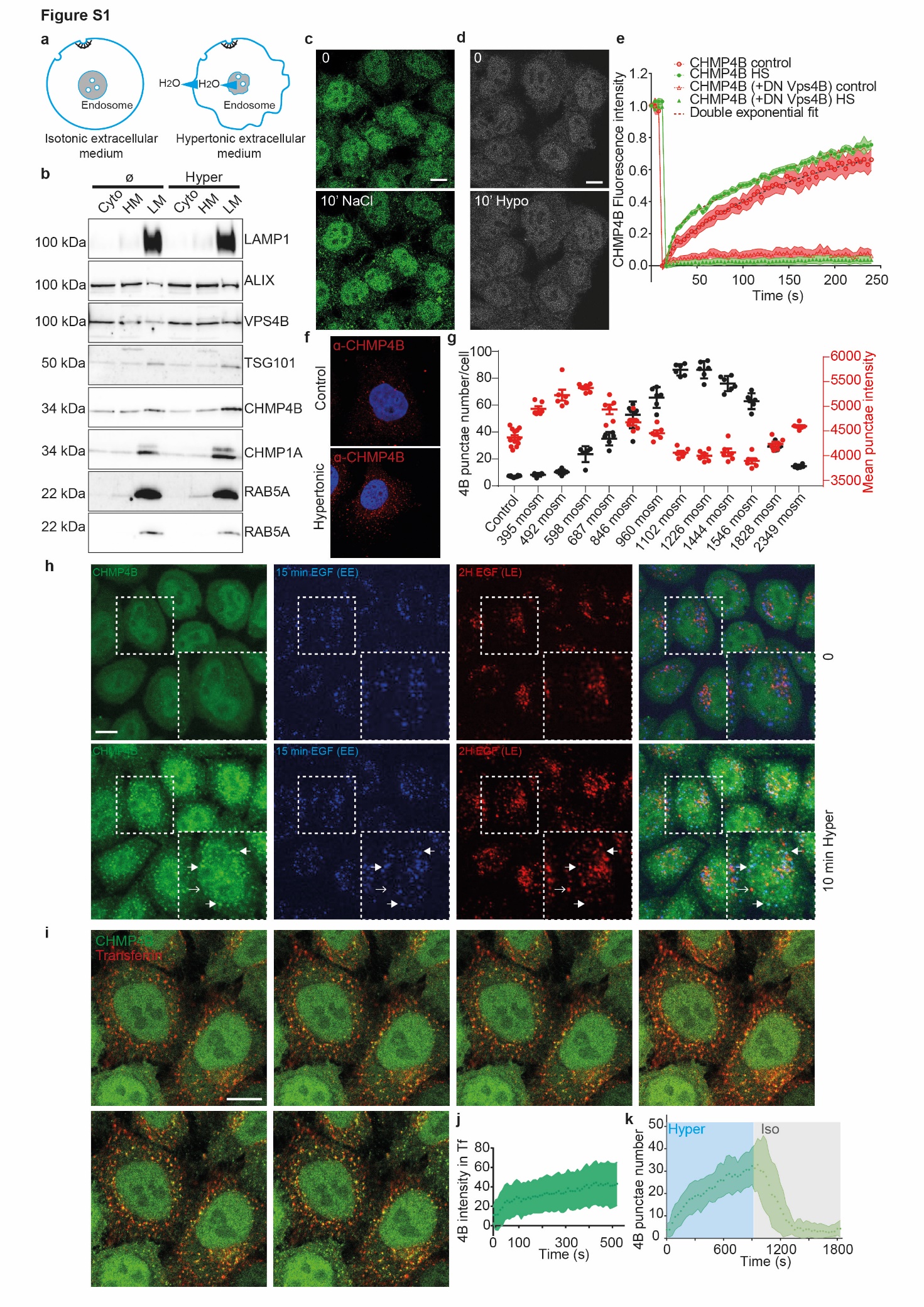

**Fig.** **S1.**

**Hypertonic shock causes relocalization of ESCRT III to early endosomes**

a) Scheme of the effect of hypertonic conditions on the cellular and endosomal volumes and membranes. b) Western blot analysis for several ESCRT and endosomal markers of light-membrane (LM), heavy-membrane (HM) and cytosol (CYTO) fractions in control or hypertonic treated cells (830 mOsm). c-d) Representative confocal images of Hela cells expressing CHMP4B-GFP before and 10min after incubation with medium containing 0.25M NaCl (c) or hypotonic medium (d). e) Fluorescence recovery after photobleaching curves of CHMP4B-GFP fluorescence on endosomes under control and hypertonic conditions (830 mOsm), in Hela CHMP4B-GFP cells transfected or not the dominant-negative mutant of VPS4B (DN VPS4B) were incubated with an isotonic (330 mOsm) or hypertonic (830 mOsm) solution for 5min. Shaded areas correspond to the SEM. f) Immunofluorescence confocal images of Hela MZ cells stained for endogenous CHMP4B after incubated 10min incubation in isotonic or hypertonic (~830 mOsm). g) Number per cell (black) and intensity (red) of CHMP4B-GFP punctae after 10min incubation of hypertonic shock at different osmolarities (N±SEM, >2000 cells quantified per condition). h) Representative confocal images of Hela CHMP4B-GFP loaded with EGF-Alexa 647 for 15min (blue, EE) and for 15min followed by a 2h chase with EGF-Alexa 594 (red, LE) under control condition (upper panel) or after 10min hypertonic shock (bottom panel). i) Representative confocal images of Hela-CHMP4B-GFP cells labelled 10min with Transferrin-Alexa594 and subjected to hypertonic shock. j) Quantification of CHMP4B-GFP fluorescence intensity within the Tf-A594 mask in the experiment shown in H. Shaded area represent SD. k) Number of CHMP4B-GFP punctae with time during a 15 min hypertonic shock (~830 mOsm), followed by a 15min isotonic recovery (~330 mOsm). Shaded areas: mean ± SD (N=27-37 cells from 3 independent replicates).

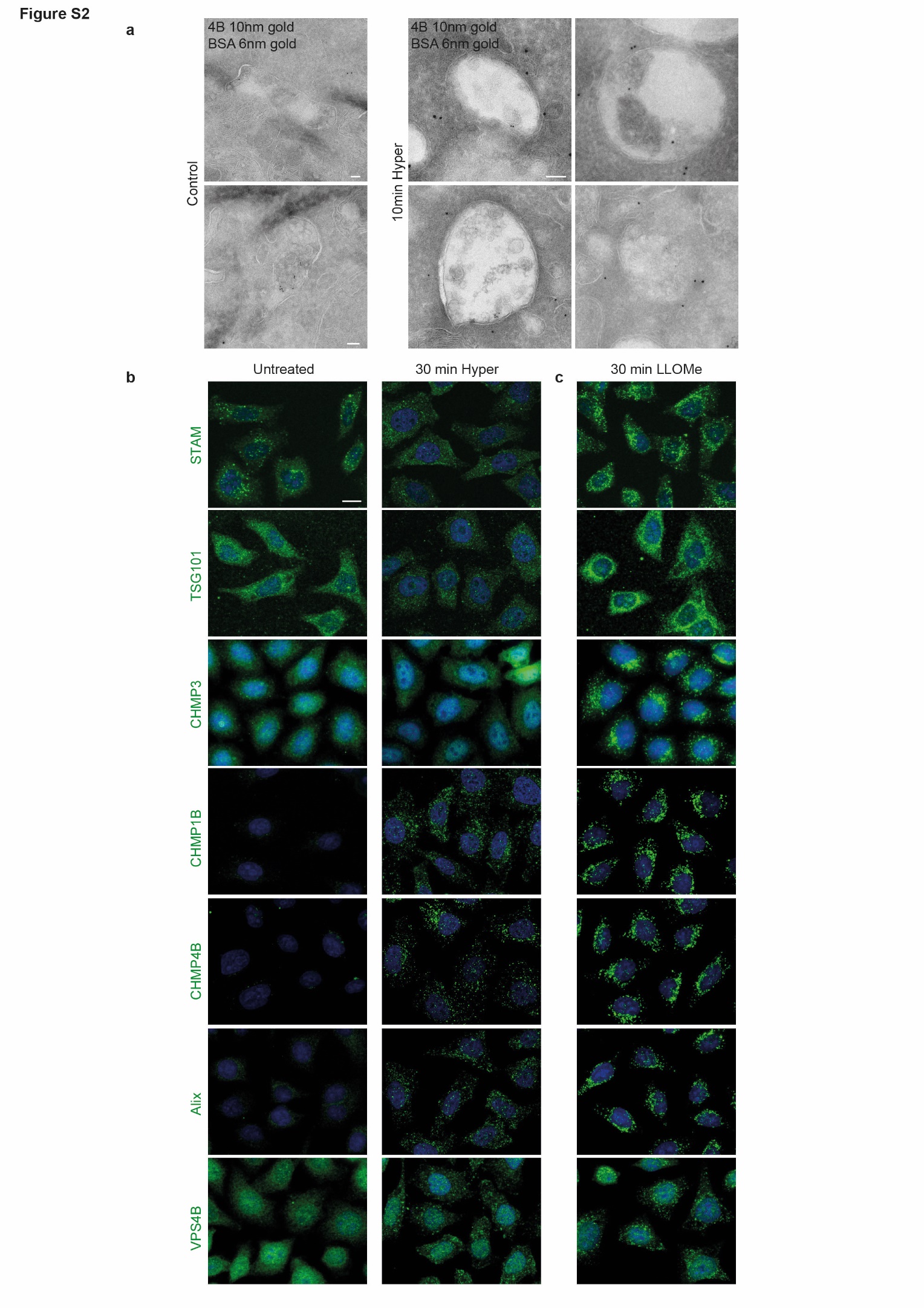

**Fig. S2.**

**Hypertonic shock or LLOMe treatment leads to ESCRT proteins relocalization**

a) Hela-CHMP4B-GFP cells were incubated for 15min with 6nm BSA-Gold and subjected to a hypertonic shock for 10min. and cells were processed for immune-electron microscopy, as in Fig 1k.The gallery shows representatives micrographs of endosomes after immuno-gold labelling of cryo-sections with anti-CHMP4B antibodies followed by 10 nm Gold-Protein A. b-c) Representatives automated confocal images of Hela MZ cells after immunostaining of the indicated ESCRT subunits under the following conditions: (b) untreated and 30min hypertonic shock, (c) 30 minutes treatment with LLOMe. Quantifications in Fig 1L and Fig 2N were obtained from similar images. Scale bar: 10 μm

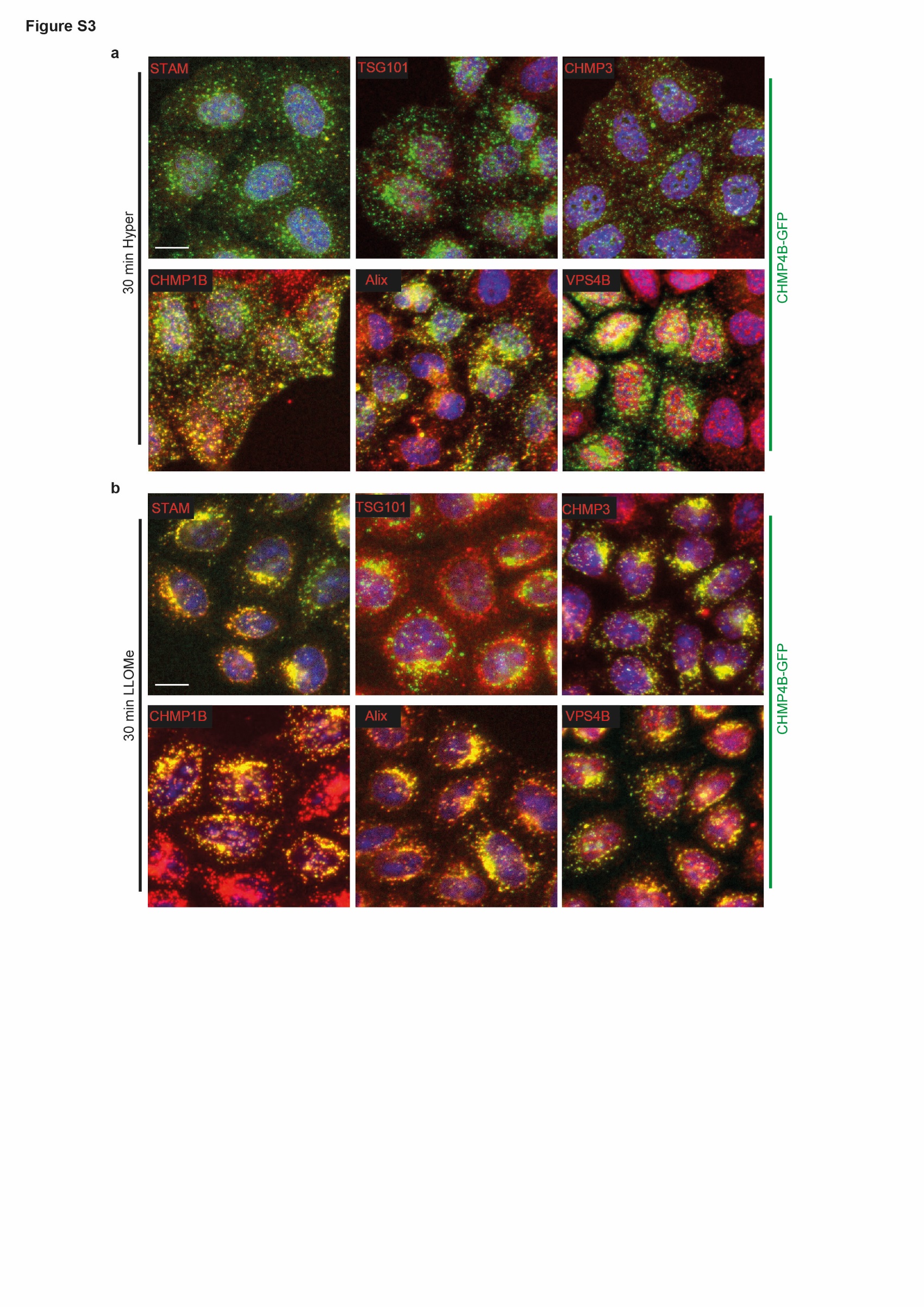

Fig. S3.

**ESCRT proteins relocalization upon hypertonic or LLOMe treatments.**

Representatives automated confocal images of Hela cells expressing CHMP4B-GFP (green) after hypertonic shock for 30min (a) or LLOMe treatment for 30min (b). Cells were labelled with antibodies against the indicated ESCRT subunits (red). Scale bar: 10 μm

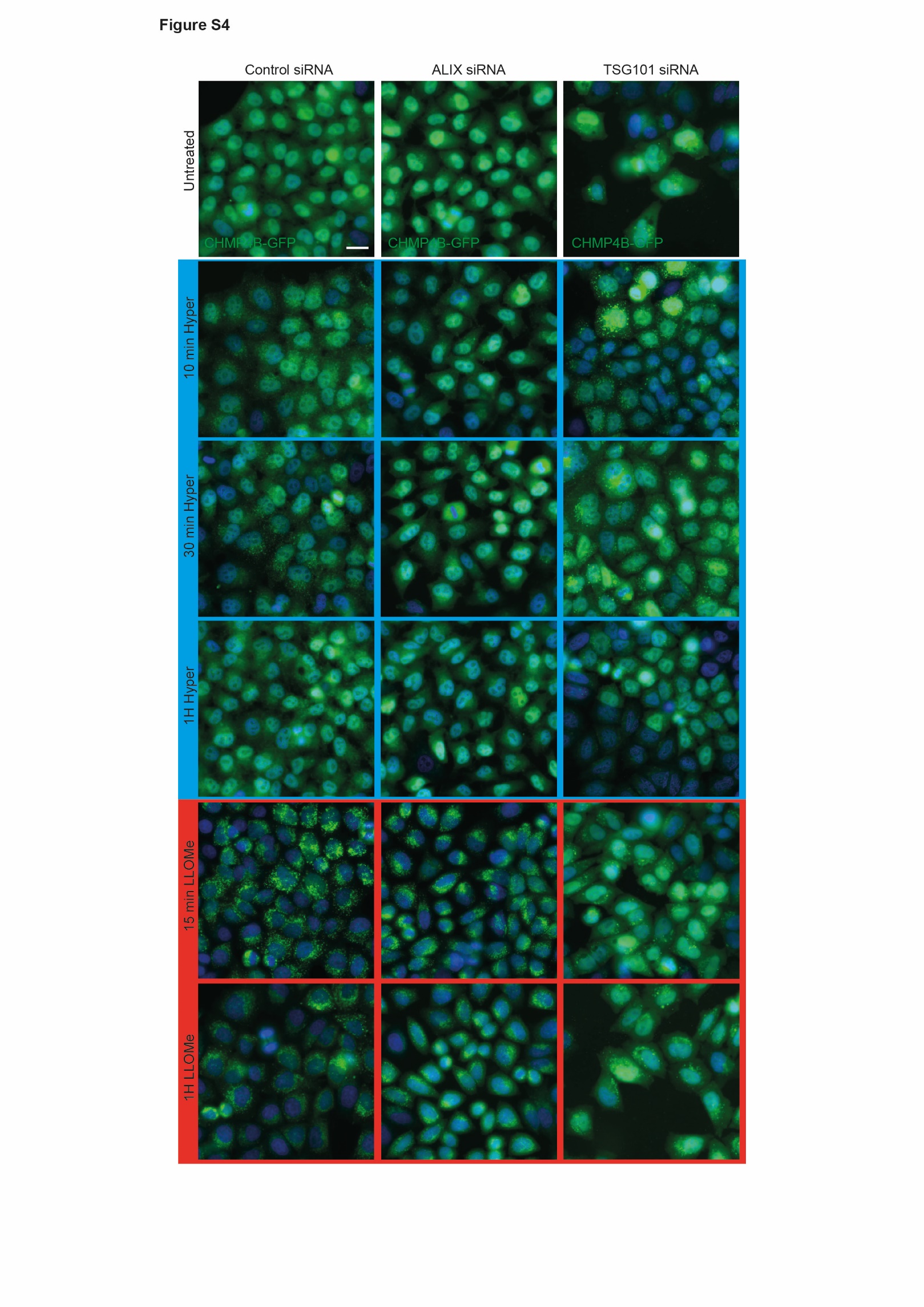

Fig. S4.

**Role of ALIX and TSG101 in the redistribution of CHMP4B after hypertonic shock and LLOMe treatment.**

Cells expressing CHMP4B-GFP were transfected with control siRNAs (siVSV) or with siRNAs against ALIX or TSG101, treated with hypertonic medium (0.5 M sucrose) or with 0.5mM LLOMe for the indicated time, and then imaged by automated confocal microscopy. The micrographs represent CHMP4B-GFP distribution and correspond to the quantification shown in Fig2q and FigS6f. Bar: 20 μm

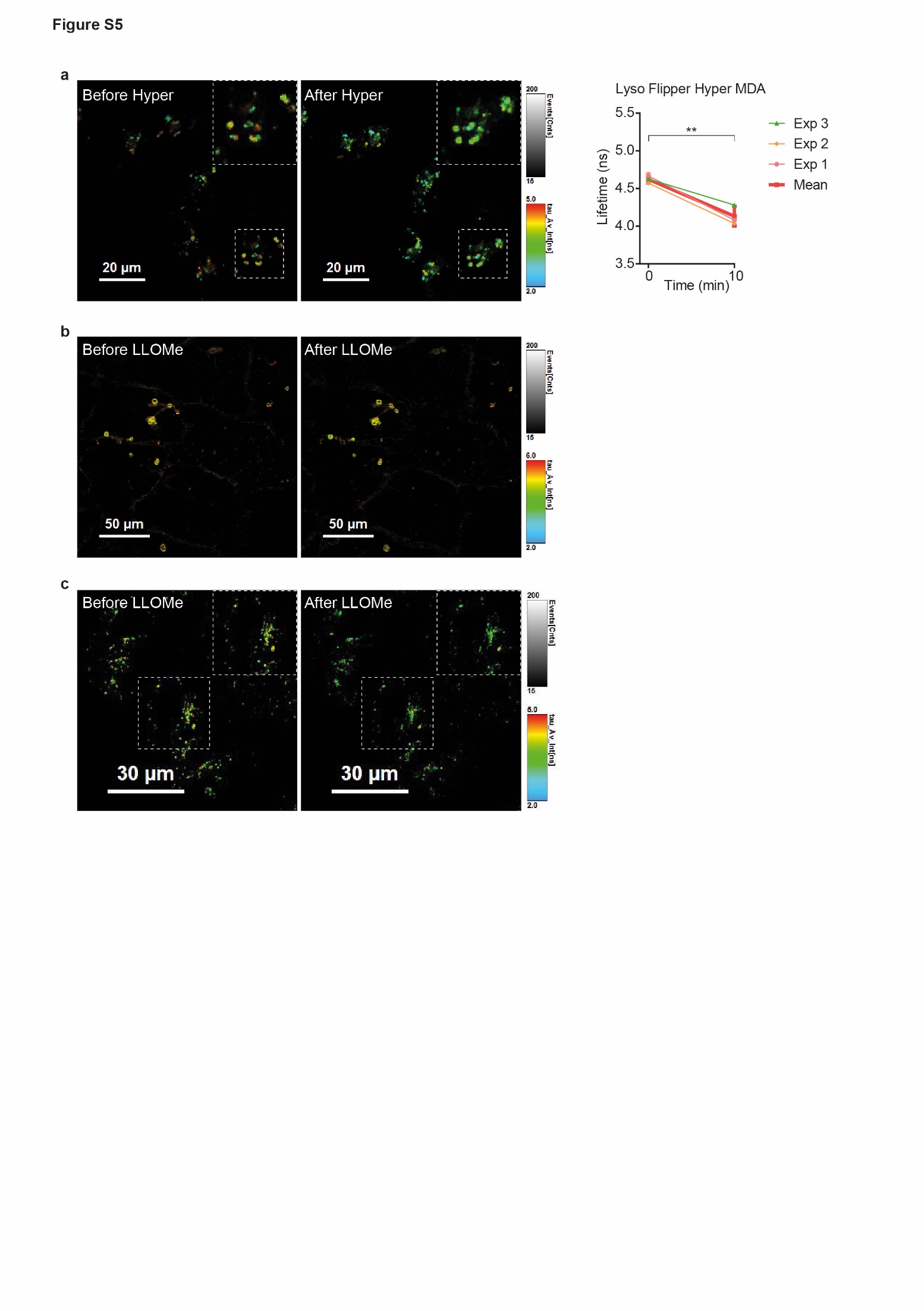

Fig.S5.

**Hypertonic shock or LLOMe treatment decrease membrane tension of endosomes**

a) Representative FLIM images of endosomes in MDA cells stained with Lyso-Flipper before and after hypertonic shock. The graph shows the quantification of these experiments where each thin line is an individual experiment (with at least 3 cells and several tens of endosomes) and the thick line the mean of these 3 experiments. Error bars represent SEM. (two-tailed paired t-test: P=0.00369362). b) Representative FLIM images of RAB5Q79L endosomes in Hela MZ cells stained after a 2h incubation with FliptR before and after 0.5 mM LLOMe treatment. c) Representative FLIM images of endosomes in Hela MZ cells stained by Lyso Flipper before and after 0.5 mM LLOMe treatment.

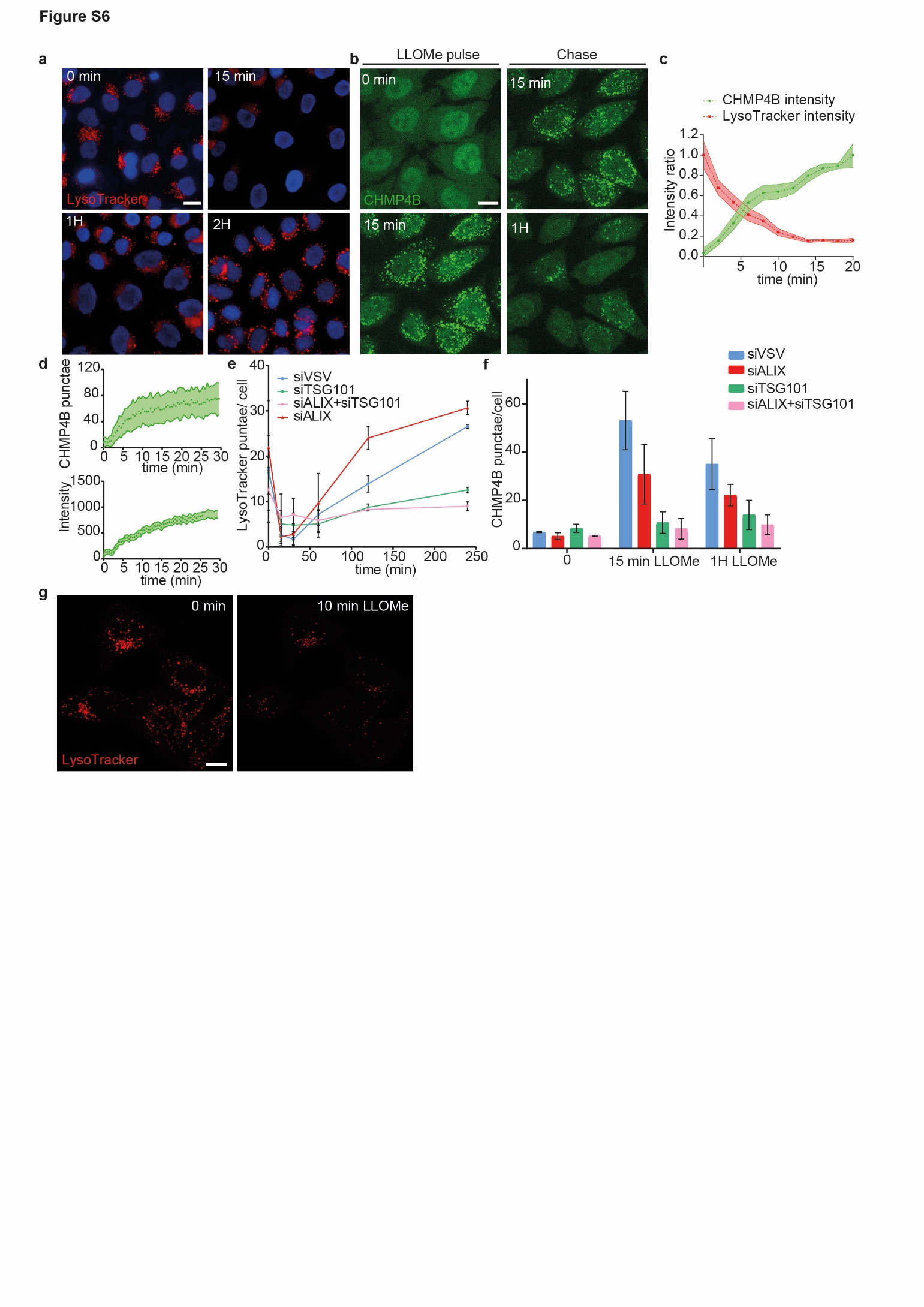

Fig. S6.

**ESCRT are recruited to late endosomes after membrane injury and necessary for their repair.**

a) Hela cells were treated for the indicated time with 0.5mM LLOMe, stained with LysoTrackerLysoTracker and imaged by automated confocal microscopy. Scale bar: 10 μm. b) Representatives confocal images of CHMP4B-GFP distribution during a pulse-chase experiment with 0.5 mM LLOMe. Scale bar: 10μm. c) The graph shows the normalized fluorescence intensity of LysoTracker and CHMP4B-GFP extracted from the Movie 3 that illustrates the treatment of HeLa cells with 0.5mM LLOMe. Shaded areas correspond to SEM from 26 ROIs in 10 cells from 2 independents movies. d) The graphs show the number of CHMP4B-GFP punctae per cell in cells treated with 0.5 mM LLOMe (upper graph, shaded area represents SD, N= 8 cells ), as well as the mean intensity of the punctae (shaded area represents SEM, N=43 ROIs in 8 cells). e) Cells expressing CHMP4B-GFP were transfected with control siRNA (siVSV) or siRNAs against ALIX, TSG101 or both ALIX and TSG101, and treated with 0.5 mM LLOMe for the indicated time. Acidic compartments were revealed using LysoTrackerLysoTracker and imaged by automated confocal microscopy. The graph shows the mean number of LysoTrackerLysoTracker punctae per cell. Error bars represent SEM (>1000 cells in 3 independent replicates). f) Cells expressing CHMP4B-GFP were transfected with control siRNA (siVSV) or siRNAs against ALIX, TSG101 or both ALIX and TSG101, and treated with 0.5 mM LLOMe for the indicated time. The graph shows the mean number of CHMP4B-GFP punctae per cell. Error bars represent SEM (>1000 cells in 3 independent replicates). g) Confocal images of LysoTracker staining before and 10min after 0.5 mM LLOMe treatment. The data correspond to Fig2I. Bar: 10 μm

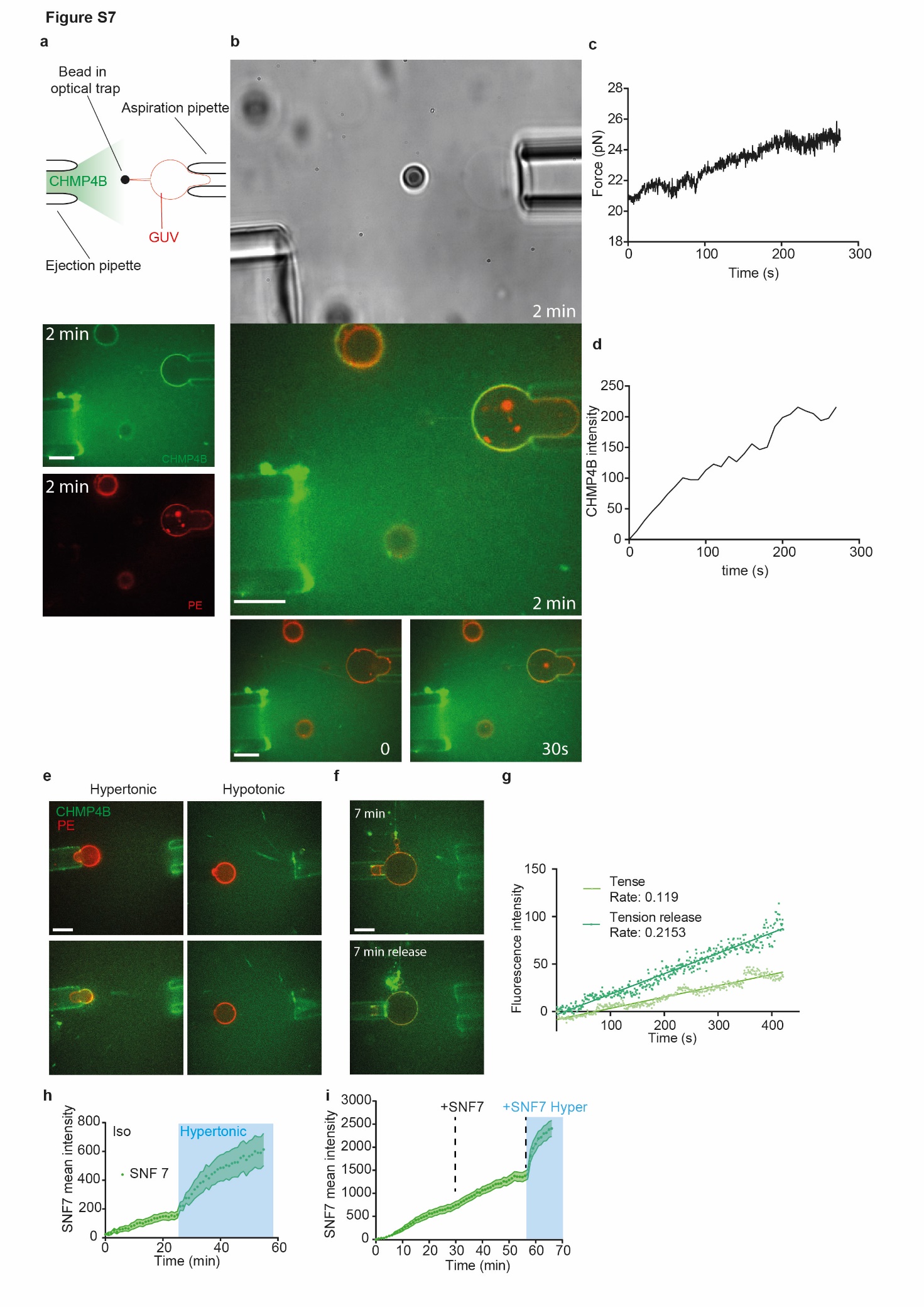

**Fig. S7.**

**Low membrane tension facilitates CHMP4B association with GUV membrane.**

a) Schematic representation of the tube pulling setup used in this study. b) Representative images of a GUV held in the aspiration pipette while purified CHMP4B-A488 is released from the ejection pipette. Binding is clearly visible 2min after CHMP4B ejection. Bars: 10 μm. c-d) The force exerted on the bead (measured with optical tweezers) increases with time (c), as does CHMP4B binding on the GUV (d). Hence, the force increase is correlated with CHMP4B binding. e) Representative images of CHMP4B-A488 binding on the GUV surface when the buffer containing CHMP4B is hypertonic or hypotonic. Bar: 10 μm. f-g) Tension was released by a decrease in the aspiration of the tongue with the aspiration pipette. The upper image (f) shows the CHMP4B fluorescence on the GUV 7min after the beginning of protein ejection and the bottom image (f) 7min after the release of membrane tension (note the disappearance of the tongue. Bar: 10 μm. Quantification of this experiment is shown in (g). h) Same experiments than in Fig 3c but with Snf7 instead of CHMP4B (at the same concentration). Error bars represent SEM (N=13 GUVs) i) GUVs were incubated for 30min with 1µm Snf7 in isotonic buffer, and then the solution was replaced with exactly the same solution (+SNF7). Then, at 56min, the solution was replaced again but with hypertonic buffer also containing 1µm Snf7 (+SNF7 Hyper, blue part of the graph). Error bars represent SEM (N=20 GUVs).

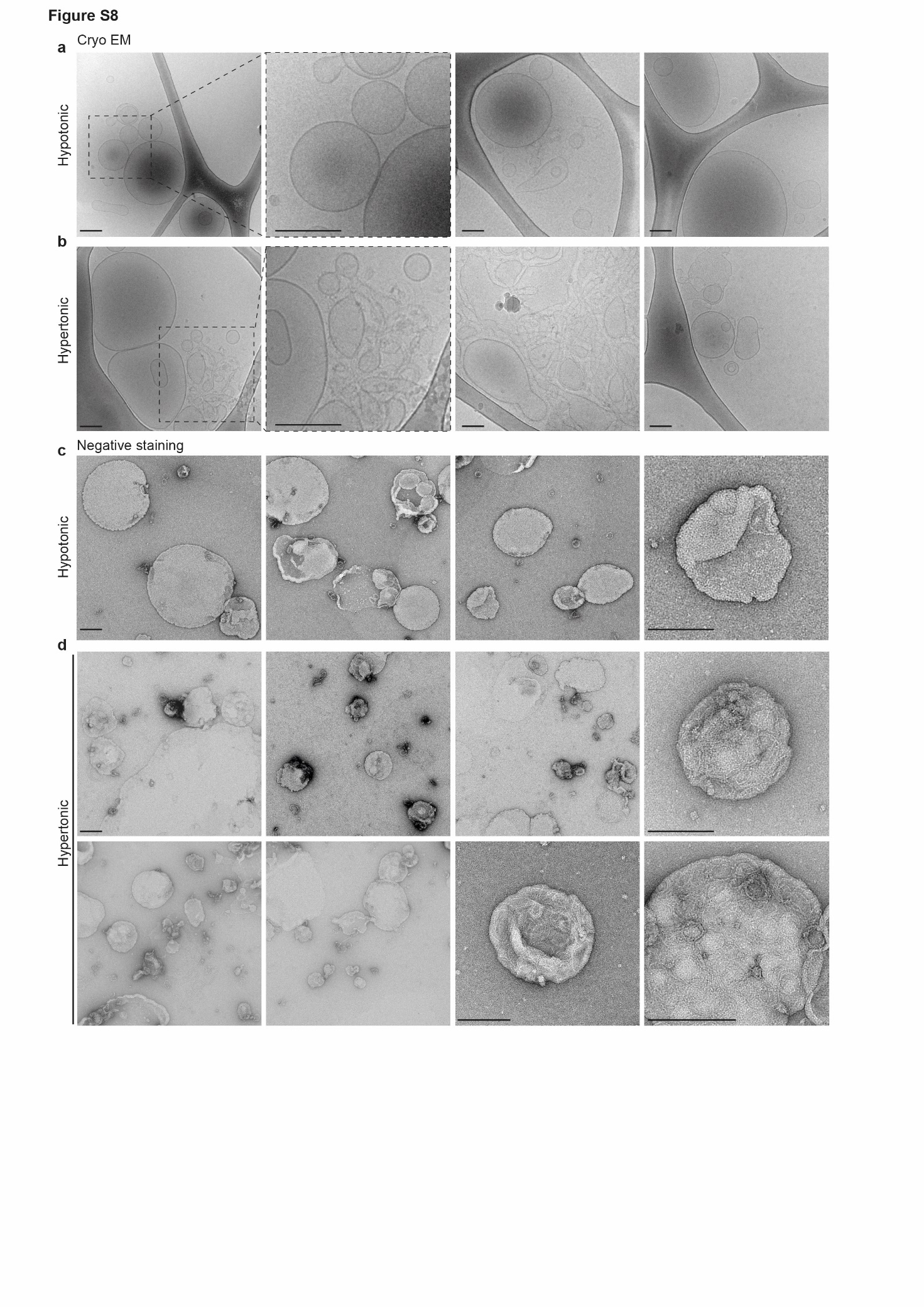

Fig. S8.

Low membrane tension promotes CHMP4B polymerization on LUVs membrane and EGF triggers endosomal membrane tension decrease in vivo.

a, b) Cryo-electron micrographs of LUVs incubated for 2h with 1μM CHMP4B-GFP in hypotonic (a) or hypertonic (b) solution. Bars: 100 nm. c, d) Electron micrographs after negative staining of LUVs incubated for 2h with 1μM CHMP4B-GFP in hypotonic (c) or hypertonic (d) solution. Bars: 100 nm.

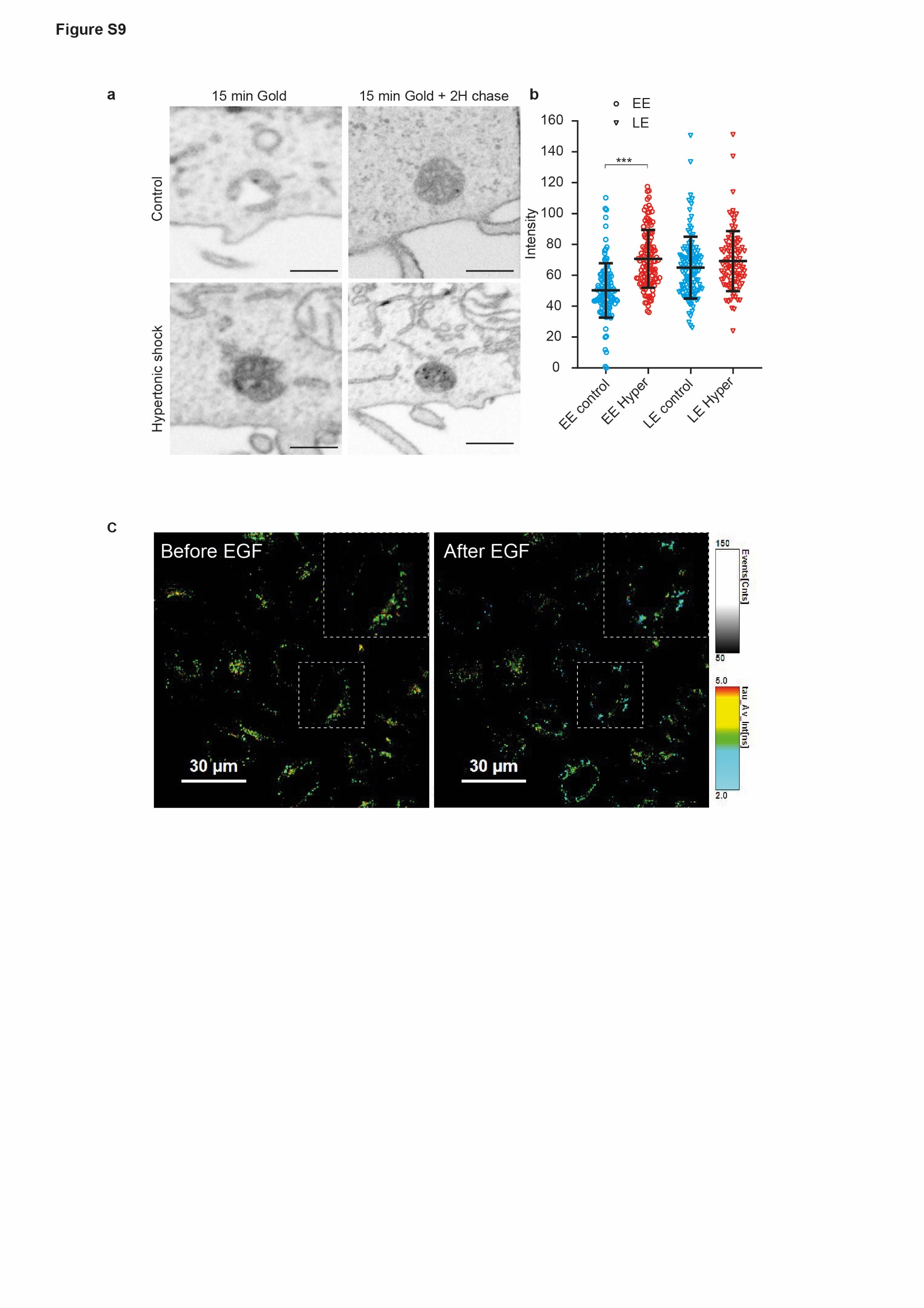

Fig. S9.

Low membrane tension promotes CHMP4B polymerization on LUVs membrane and EGF triggers endosomal membrane tension decrease in vivo.

a-b) As in Fig 4, the panel (a) shows FIB-SEM micrographs of cells loaded with BSA-gold. The electron density of BSA-gold labelled early endosomes (EE), or late endosomes (LE) was measured before (blue) and after (red) a 10min hypertonic shock (b). Error bar is SD (N: EE control:121; EE Hyper: 124; LE control:120; LE Hyper:98 in 2 independent replicates, Kruskal-Wallis test: P<1.10^-15^). c) Representative FLIM images of Hela MZ endosomes stained with Lyso-Flipper before and after 20min 200 ng/ml EGF treatment.

Movie S1.

Hypertonic shock triggers fast relocalization of CHMP4B to endosomes

Confocal time-lapse movie recorded at speed of 1 frame/10sec of CHMP4B-GFP during hypertonic treatment.

Movie S2.

Hypertonic shock triggers transient relocalization of CHMP4B to endosomes

Confocal time-lapse movie recorded at speed of 1 frame/5 minutes of CHMP4B-GFP during hypertonic treatment. This movie corresponds to the time series shown in Fig. 1A.

**Movie S3**

**LLOMe triggers massive relocalization of CHMP4B.**

Confocal time-lapse movie recorded at speed of 1 frame/30sec of CHMP4B-GFP during 0.5 mM LLOMe treatment.

Movie S4.

LLOMe triggers loss of endosomal acidity and quick relocalization of CHMP4B to endosomes

Confocal time-lapse movie recorded at speed of 1 frame/2min of CHMP4B-GFP cells stained with LysoTracker during 0.5 mM LLOMe treatment.

Movie S5.

A decrease in membrane tension trigger by hypertonic buffer increases CHMP4B polymerisation rate in vitro

Confocal time-lapse movie recorded at speed of 1 frame/min of CHMP4B-A488 on GUVs during isotonic followed by hypertonic incubation.
